# Teacher-student collaborated multiple instance learning for pan-cancer PDL1 expression prediction from histopathology slides

**DOI:** 10.1101/2023.07.26.550748

**Authors:** Darui Jin, Shangying Liang, Artem Shmatko, Alexander Arnold, David Horst, Thomas G. P. Grünewald, Moritz Gerstung, Xiangzhi Bai

## Abstract

Programmed cell death ligand 1 (PDL1), as an important biomarker, is quantified by immunohistochemistry with few established histopathological patterns. Deep learning aids in histopathological assessment, yet heterogeneity and lacking spatially resolved annotations challenge precise analysis. Here, we present a weakly supervised learning approach using bulk RNA sequencing for PDL1 expression prediction from hematoxylin and eosin (H&E) slides. Our methods, MILTS, extends multiple instance learning paradigm with the teacher-student framework, which assigns dynamic pseudo-labels for intra-slide heterogeneity and retrieves unlabeled instances using temporal ensemble model distillation. The approach, evaluated on 12,299 slides across 20 solid tumor types, achieves a weighted average AUC of 0.83 on fresh-frozen and 0.74 on formalin-fixed specimens for 9 tumors with PDL1 as an established biomarker. MILTS predicts PDL1 expression patterns, validated by immunohistochemistry on 20 slides, offering insights into histologies relevant to PDL1. This demonstrates the potential of deep learning in identifying diverse histological patterns for molecular changes from H&E images.

## Introduction

Inhibitors for the PD1-PDL1 checkpoint have revolutionized cancer therapy in the past decade. In addition to seven anti-PD1/PDL1 monoclonal antibodies (mAbs) currently approved by US Food and Drug Administration (FDA), there are still approximately six thousand mAbs undergoing clinical trials [1–3]. Blockade of PD1-PDL1 interaction notably contributes to activating antitumor immunity and is proven to benefit the treatment of various types of tumors [4–6]. PDL1 expression serves as a biomarker associated with patients’ response to anti-PD1/PDL1 mAbs such as pembrolizumab and nivolumab, which acts as the most widely adopted standard for identifying patient cohorts that are appropriate candidates for PD1-PDL1 immunotherapy [7]. From 2011 to 2021, there were totally 15 immune checkpoint inhibitor FDA approvals linked with companion PDL1 testing including non-small cell lung cancer (NSCLC) (N=7), bladder cancer (N=3), triple-negative breast cancer (N=2), cervical cancer (N=2) and gastric cancer (N=1). And the outcomes of patients with renal cell cancer, colon cancer and melanoma are also reported linked with PDL1 expression by many researchers [8–12].

Currently, PDL1 expression is predominantly quantified by immunohistochemistry (IHC) assays, and some recent research has also indicated a significant correlation between mRNA expression levels and the response of associated monotherapies[13]. Whilst IHC qualification has been successfully used in clinical practice for decades and is considered to be a gold standard for this task, recent analysis show that the interpretation of staining and decision threshold differs for different commercially available platforms and even differs within the same platform [14]. Along with the subjectivity of pathologists, these factors introduce undesired inter– and intraobserver variance to the evaluation of staining and thus limited reproducibility. Besides, quantification of mRNA expression using techniques like real-time reverse transcription polymerase chain reaction (RT-qPCR) and IHC tests can be costly and time-consuming[15]. H&E stained slides are one of the most widely used and effective carriers of pathological information, offering a cost-effective and expedient alternative, and are employed in routine pathological assessment of clinical specimens. Developing an efficient and reliable method for estimating PDL1 expression on H&E stained slides, which does not require additional sample preparations, may yield a faster and cheaper diagnostic readout. Further, this process is capable of unraveling histopathological characteristics of PDL1 expression on slides, thereby assisting pathologists in comprehending the gene’s expression mechanism and facilitating a more precise interpretation of immune evasion patterns.

The expanding field of computational histopathology may not only automate existing workflows, but also proposes to explore the molecular information based on morphological features via deep learning [16]. It has exhibited great potential in assisting pathologists on many routine tasks [17] including applications such as mitosis detection [18–21], tissues segmentation [22–25], tumor subtyping and grading [26–29] and biomarker assessment [30–33]. Some studies also reveal that morphotypes are closely associated with genetic alterations in tumors and thus indicative of clinical features and prognosis, which has a chance to redefine the clinical workflows [34–37]. However, the analysis of pathological images is often challenged by the limited availability of reference data, with clinical reports and patient-level diagnoses being the main sources of information[16]. And it is also tricky to accurately capture and spatially resolve transcriptome profiles at the pixel or cell level in whole slide images [38]. The presence of such heterogeneity poses challenges in training deep learning based algorithms, which usually requires accurate reference in the form of labeled data[39]. Some studies have resorted to manual data annotation and adopted fully supervised learning strategies. For example, in the works of Sha et al. [40] and Shamai et al. [33], PDL1 expression quantification was performed at the tile level by pathologists using paired IHC slides providing an accurate reference for training. However, the dataset size could be constrained due to the labor-intensive nature of annotating such data. Some existing approaches have opted to overlook the intra-tumor heterogeneity by assigning slide-level annotations to all content within a slide [31, 34, 35, 41, 42]. This strategy works well if the slides exhibit good homogeneity concerning specific properties of interest. However, it can also lead to overfitting and undesired generalization with heterogeneous composition, where inaccurate instance-level labels will either hinder the convergence of the model or result in erroneous recognition of relevant patterns [43]. More recent approaches tend to directly utilize ImageNet [44] pretrained features and incorporate specially designed attention or embedding-based MIL strategies [19, 24, 29, 36, 45–48], which significantly accelerates the training but is also more data-hungry. In addition, dimensionality reduction with pretrained features inevitably leads to information loss. These pre-trained models based on the ImageNet database, are designed to capture general visual patterns without any specific bias towards histopathology-related features or priors. Consequently, in situations where certain details in histopathology images are completely lost or unavailable, reweighting or attention mechanisms have limited effectiveness.

In this work, to leverage the benefit of both massive amount of tile information and slide-level annotation, we propose our weakly supervised learning based methodology named MILTS (Teacher-Student collaborated Multiple Instance Learning framework), which is powered by whole slide images in H&E staining from The Cancer Genome Atlas (TCGA) and The National Cancer Institute’s Clinical Proteomic Tumor Analysis Consortium (CPTAC) involving 12,299 slides from 6,715 patients across 20 kinds of tumors. MILTS utilizes an iterative, self-refining process to assign labels automatically supporting tile feature extractor training, and combines the statistical summary of tile-level predictions with tokens fused by the transformer to obtain the slide-level embedding for the final prediction. Results demonstrate there exists a salient morpho-transcriptomic link across certain cancer types, whose treatment landscape and prognosis are associated with PDL1 expression, with a weighted average AUC of 0.83 on fresh-frozen and 0.74 on FFPE specimens. Heterogeneous tile-level predictions further provide insights into morphotypes associated with PDL1 hot regions in colon cancer, which include mixed inflammatory stroma with relatively high abundance of eosinophils and a cribriform growth pattern of tumor cells with hyperchromatic nuclei. And it is observed that the model predictions consistently exhibited a strong positive correlation with corresponding IHC quantification, providing further validation for the findings based on H&E staining. Building on the foundation of PDL1-relevant tumors, we undertake an analysis of the other 11 cancer entities on the morpho-transcriptomic correlation that have not been verified or approved yet. Varying degrees of morpho-transcriptomic correlation is observed among these tumors. Further examination may be warranted based on the difference between predictability of these tumors and known PDL1-relevant ones. It also reveals that the molecular basis of tumors could be depicted from the view of cellular morphology via advanced deep learning techniques, which could provide a new perspective on studies of tumorigenesis and treatment.

## Results

### Workflow of MILTS

The corresponding workflow of MILTS is presented in Fig. 1a. In the context of MILTS, a slide-level label initialized from dichotomised mRNA expression levels is employed to supervise the representation learning for the histopathological image, where three quantiles (quartile, tertile, and median points) for each cancer type were considered. Specific values could be found in Supplementary Tables S1 and S2. Because the slide-level labels constitute an aggregate summary, which is expected to differ across the tiles of the tumor section, the teacher-student framework combines dynamic labels assignment for individual tiles with knowledge distillation from a temporal ensemble model representing the exponentially decaying average of previous learning iterations. Specifically, tiles would be fed into both the teacher and student model with random augmentation including rotation, crop, flip and color transformation. The teacher model continuously yields tile-level pseudo-labels for typical positive/negative tiles in each epoch, based on which the student model would be updated following the MIL constraint as well as the distribution generated by the teacher model on unlabeled instances. As the teacher model is continuously updated via the moving average of the student model, this collaborative procedure automatically learns optimal tile level labels. These features are further fused by a transformer to obtain a slide-level token and combined with statistical summary of patch-level outcomes in multi-layer perceptron (MLP) to infer per patient results. More details about the methodology are provided in the Methods section.

**Fig. 1.**
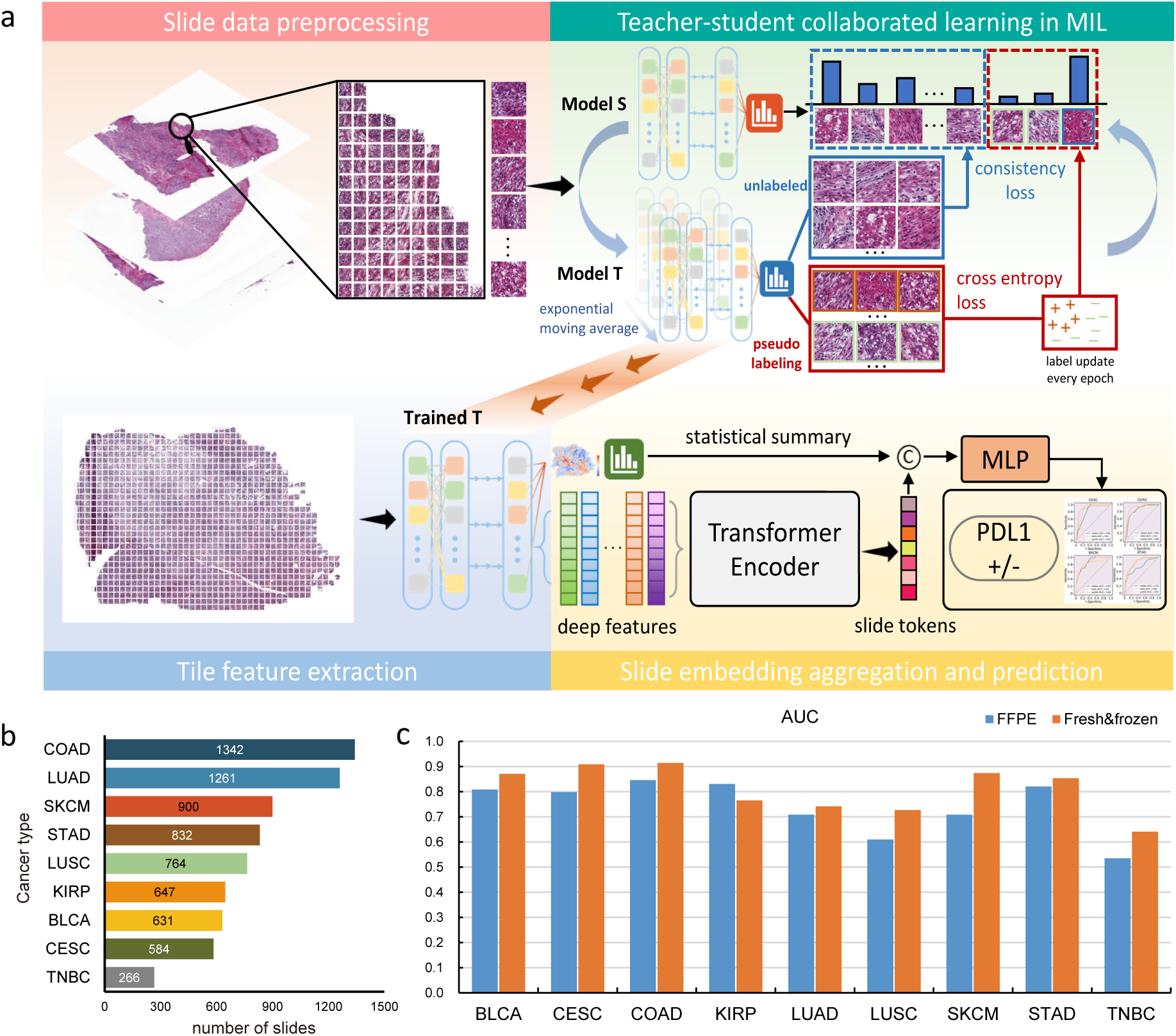
The framework of MILTS and its performance on clinically PDL1-relevant tumors. **a**, The training and inference workflow of MILTS. The training mainly includes three steps. First the data of patient cohorts are divided into training set, validation set and test set, which would be followed by patching and random augmentations. Then these tiles are utilized to train the patch-level teacher-student collaborated network in a multiple instance learning manner. At last, the trained patch-level teacher model (or student model) works as the extractor of both statistical features and deep features. The deep features of patches in the same slide are further fused into a slide token and combined with the statistical summary of patch-level features to train an MLP classifier which would give the patient-level diagnosis. MIL, multiple instance learning; S, student; T, teacher; C, concatenation; MLP, multi-layer perceptron. **b,** Quantities of slide images of different tumors. **c,** The plot illustrating the model’s performance on FFPE slides and fresh-frozen slides for the afore-mentioned tumors, separately. Source data are provided as a Source Data file.

### Predictability of PDL1 expression across 9 cancer types

The ability to classify gene expression was evaluated on 9 cancers for which PDL1 expression serves as an established biomarker for checkpoint inhibitors, whose distribution is shown in Fig. 1b (involving 3,121 cases with 4,215 fresh-frozen slides and 2,966 FFPE slides). Using the upper tertile of PDL1 expression as a threshold, MILTS achieved performance on fresh-frozen slides with a weighted average average area under receiver operating characteristic curve (AUC) of 0.83 (range: 0.64 to 0.91), accuracy of 0.75 (range: 0.58 to 0.87), sensitivity of 0.83 (range: 0.74 to 0.90) and specificity of 0.71 (range: 0.47 to 0.89). We also evaluated the using other two threshold settings: the upper quartile (top 75%) and median (50%) expression levels in each cancer type. These two alternative thresholds yielded broadly comparable performance with a mean AUC of 0.81 and 0.75, respectively (Supplementary Fig. S3 and Supplementary Tables S3 and S4). In the following, we discuss results at the upper tertile, unless stated otherwise.

Specifically, the dataset comprises bladder urothelial carcinoma (BLCA), cervical squamous cell and endocervical adenocarcinoma (CESC), colon adenocarcinoma (COAD), kidney renal papillary cell carcinoma (KIRP), lung adenocarcinoma (LUAD), lung squamous cell carcinoma (LUSC), skin cutaneous melanoma (SKCM), stomach adenocarcinoma (STAD) and triple-negative breast carcinoma (TNBC). The results are shown in Fig. 1c. To benchmark performance across various tumors, the threshold to determine the accuracy, sensitivity and specificity is selected by Youden’s J statistic. The optimal threshold here is the one that maximizes the sum of sensitivity and specificity. This strategy is applied across all our experiments unless stated otherwise. 95% confidence interval (CI) is also computed with bootstrapping strategy (2,000 random resamples), where the AUC performances on fresh-frozen slides are respectively 0.87 for bladder urothelial carcinoma (95% CI: 0.84-0.90), 0.91 for cervical squamous cell and endocervical adenocarcinoma (95% CI: 0.88-0.93), 0.92 for colon adenocarcinoma (95% CI: 0.90-0.93), 0.77 for kidney renal papillary cell carcinoma (95% CI: 0.73-0.80), 0.74 for lung adenocarcinoma (95% CI: 0.72-0.77), 0.73 for lung squamous cell carcinoma (95% CI: 0.70-0.76), 0.87 for skin cutaneous melanoma (95% CI: 0.86-0.89), 0.85 for stomach adenocarcinoma (95% CI: 0.83-0.87) and 0.64 for triple-negative breast carcinoma (95% CI: 0.60-0.69).

Separate models were trained on FFPE samples using the tertile threshold. We maintained consistent cohort splits employed for fresh-frozen sections and an average AUC of 0.74 was achieved with the model trained with FFPE slides, where the trend in tumor-specific performance was consistent with that of fresh-frozen slides. Detailed results are presented in Supplementary Tables S5 and S6. Nonetheless, there remained a performance gap between FFPE slides and fresh-frozen ones as shown in Fig. 1c. The finding that frozen slides usually yield better molecular inference aligns with observations reported in several previous studies [26, 32]. Further investigations are warranted to explore strategies for bridging the gap between these two modalities. Overall, a significant morpho-transcriptomic link is evident regarding PDL1 expression across nine PDL1-relevant cancer types, which demonstrates a good predictability independent of the specific threshold.

### MILTS outperforms other methods in PDL1 expression prediction

Comparison results clearly demonstrate that MILTS outperforms the comparison methods on aforementioned tumors, exhibiting a notable advantage of 9% or more in terms of average AUC. Several methods were evaluated including the method by Campanella et al. [36], TransMIL [46] and CLAM [47]. All comparison methods are deep learning based MIL algorithms, among which Campanella’s adopts instance-level strategy while TransMIL and CLAM are embedding-level ones. The related hyperparameter settings are listed in Supplementary Table S7. Quantitative results are provided in Fig. 2a. Overall, MILTS gains superior performance to the comparison methods when making predictions about the expression level. The weighted average AUC of MILTS on these 9 kinds of tumors is 0.83, while those of the method by Campanella et al., TransMIL and CLAM are respectively 0.63 (range: 0.51 to 0.75), 0.71 (range: 0.53 to 0.84) and 0.74 (range: 0.56 to 0.83). The results of F1 score and Matthews correlation coefficient (MCC) further reveal that MILTS exhibits more balanced sensitivity to both positive and negative samples, with an average performance of 0.69 and 0.50, respectively. In comparison, the second-best model achieves only 0.60 and 0.39 for the same metrics.

**Fig. 2.**
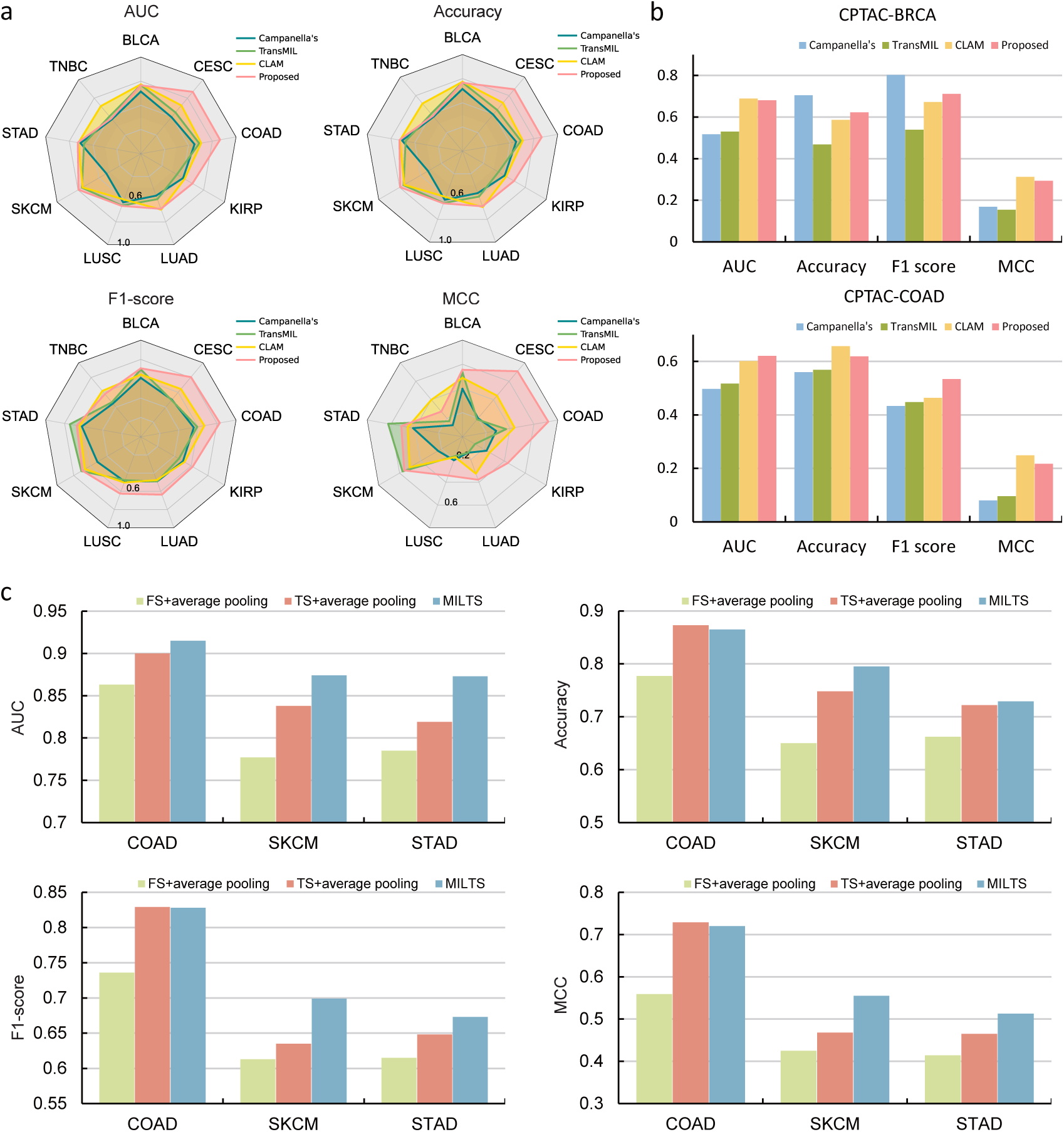
Quantitative comparison of PDL1 expression in clinically relevant tumors with other methods and ablation study results. **a**, The radar charts, arranged from left to right and top to bottom, represent the AUC, accuracy, F1 score, and MCC of the proposed method and comparison methods, respectively, at the tertile threshold. **b,** The histogram shows the results of external validation on the CPTAC BRCA and COAD datasets using the same thresholds as in the TCGA datasets. **c,** The histograms of ablation study with respect to AUC, accuracy, F1 score and MCC. In the group of ‘FS + average pooling’, a fully supervised framework and average pooling of patch-level predictions was adopted. In the group of ‘TS + average pooling’, the patch-level feature aggregation module of MILTS was substituted with average pooling. Source data are provided as a Source Data file.

An ablation study was conducted to evaluate the individual modules of the teacher-student MIL learning and feature aggregation using COAD, SKCM and STAD data. Specifically, MILTS consists of two key modules: a teacher-student MIL module for patch-level feature extraction and a transformer-based feature aggregation module for slide-level predictions. The ablation study was conducted by examining these two modules in isolation. Details about the modules are provided in the Methods section. The results are presented in Fig. 2c. Results reported that teacher-student MIL module brought an improvement of 4.4%, 8.5%, 5% and 8.8% with respect to AUC, accuracy, F1 score and MCC. The improvements of the feature aggregation module are respectively 3.5%, 1.5%, 2.9% and 4.2%, respectively. Besides, we also implemented external validation on the CPTAC dataset [49] whose results are shown in Fig. 2b. The proposed method still demonstrated superior overall performance when compared to the other methods. However, all algorithms exhibit a decrease in performance when applied to the CPTAC dataset. This decline in accuracy could be attributed to several factors, including variations in mRNA quantification and differences in the slide image modalities used in the CPTAC dataset which is a mixture of fresh frozen slides and FFPE slides. Further investigation and refinement of the algorithms may be necessary to address these issues and improve their performance on the CPTAC dataset. Nonetheless, the quantitative analysis of comparison confirms that the proposed model appear to better decipher morphological features associated with PDL1 expression from pathological patterns compared to other methods.

### Spatially heterogeneous patterns of high PDL1 expression

Like most other MIL models, MILTS calculates predictions for each individual tile as well as for the entire slide. Strikingly, tiles from samples within upper tertile of bulk PDL1 expression are predicted to exhibit a wide range of PDL1 expression, whereas tiles from the remaining samples were uniformly low (Fig. 3a). These distributions of tile-level probability accord with the assumption that there exists a substantial part of instances in positive slides which may not exhibit the same characteristics as the overall slide-level label suggests. Similar results were observed for upper quartile and median thresholds, where it appears that the range of positive probabilities for the PDL1-low class tends to shrink as the threshold value increases (Fig. S4). Conversely the range of positive probabilities became wider when the median was chosen as the threshold, indicating that the model begins to unravel nuances of very low PDL1 expression. The distinct distributions between PDL1 high and low groups explain the observed differences of AUCs (Fig. S5 and S6). Noting that the distributions show greater overlap of other thresholds reflects the lower AUCs for these cutoffs and indicates that the histopathological differences are less pronounced for median and upper quartile threshold.

**Fig. 3.**
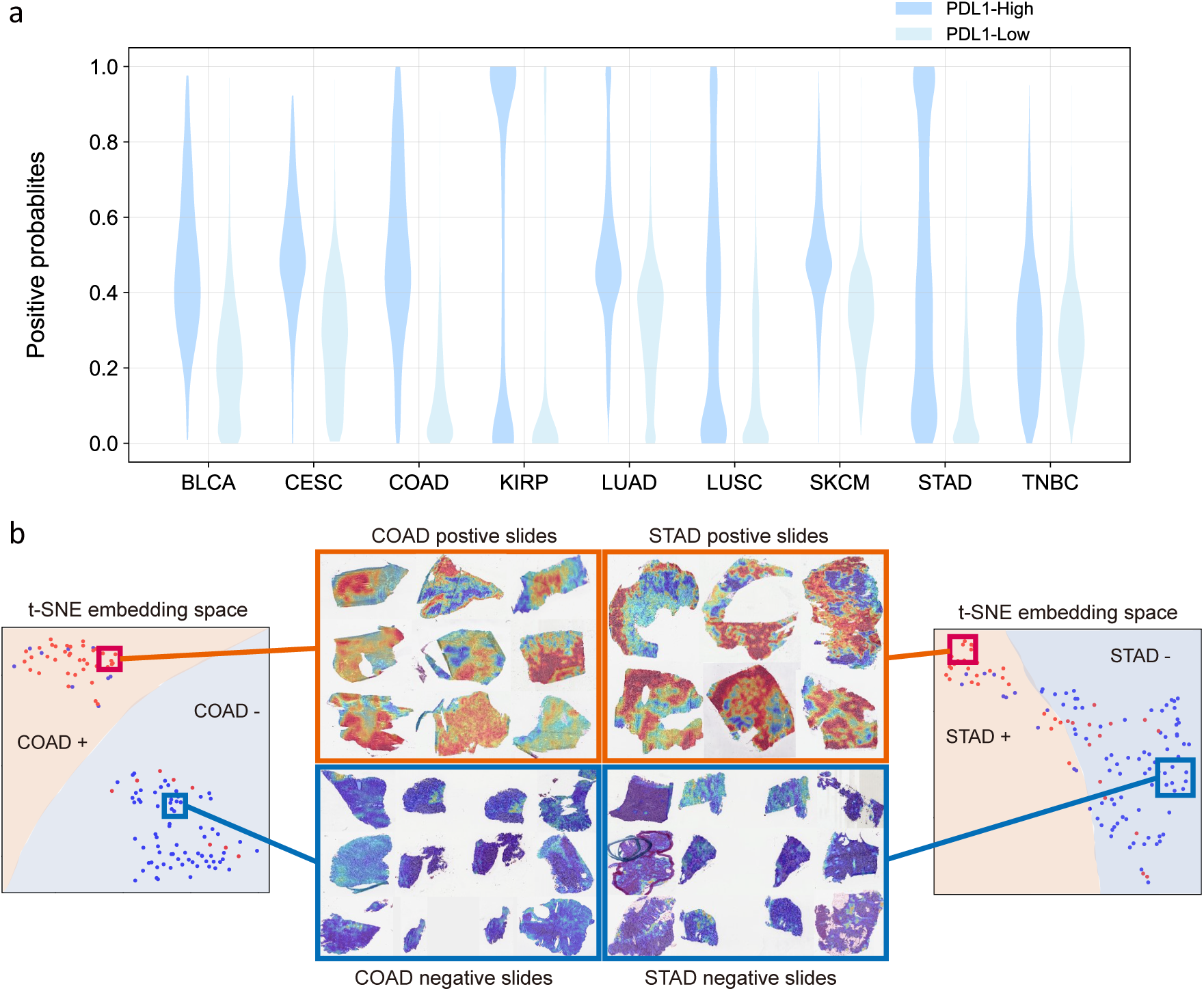
Different distributions between samples of PDL1 high and low expression. **a**, The violin plot of positive probabilities for tiles from PDL1 high class and low class at thresholds of upper tertile for each kind of tumor. **b,** Two-dimensional feature space constructed by t-SNE using 523-dimensional slide-level embedding. Heatmaps shown in the red boxes are from positive samples and those in the blue boxes are from negative samples, within which red represents a high probability of being PDL1 highly-expressed and blue indicates the opposite. Source data are provided as a Source Data file.

The notion that positive tiles reflect common histological features is also supported by a UMAP [50] visualization of the embeddings learned by the patch-level classifier in (Fig. S7). Here the representations of instances in slides with high PDL1 expression exhibited varying degrees of overlap with instances from negative slides, and those with high expression also demonstrated their own distinct distribution, independent from the distribution of negative samples, which were also reflected respectively by the cold region and highlighted region in heatmaps of PDL1 positive slides (Fig. 3b).

### Deep morphological features help discovering morphotypes for PDL1 expression

The afore-mentioned dispersion of high PDL1 predictions across tiles from the same slide manifest in a spatially organized fashion as evidenced by heatmaps of the predicted PDL1 expression (Fig. 3b). In the case of negative slides, their distribution tends to have a relatively lower mean value, resulting in a preponderance of blue regions dominating the corresponding heatmaps. In contrast, the heatmaps of positive slides exhibit different contiguous areas of predicted high and low expression values. This spatial heterogeneity also coincides with a range of histopathological patterns, which we illustrate at the example of colon adenocarcinoma.

Presented in the first row of Fig. 4a, it is common to observe a mixed inflammatory stroma characterized by a relatively high abundance of eosinophils in areas with high PDL1 expression, which can be either intact or degranulated. This suggests an immune response or inflammatory process occurring in these areas. The findings in [51] align with our observations, where PDL1 expression was predominantly observed on tumor-associated inflammatory cells by pairwise comparison with IHC slides in MSI-H subtypes. Another distinctive feature observed in areas with high PDL1 expression is the presence of a cribriform (sieve-like) growth pattern of tumor cells in the second row of Fig. 4a. This growth pattern is accompanied by tumor cells exhibiting hyperchromatic nuclei, which appear darker and more intensely stained compared to surrounding cells. The presence of cribriform growth has been identified as an independent prognostic factor in various types of cancers, indicating a higher risk of tumor progression, metastasis, and decreased overall survival. In contrast, typical negative patterns usually included areas of non-invasive adenomatous parts of the lesion as illustrated in the first row of Fig. 4b. The non-invasive adenomatous areas appear more uniform and well-differentiated glandular architecture, indicating a lower likelihood of malignancy or aggressive behavior. Other patterns also include normal colonic crypts adjacent to the invasive carcinomas, tumor necrosis, abundant tumor-associated mucin in case of mucinous carcinomas and coagulation necrosis at the sample margin in Fig. 4b. The aforementioned recurring patterns were identified in both fresh-frozen and FFPE slides. Additional examples can be found in Figures S8 and S9.

**Fig. 4.**
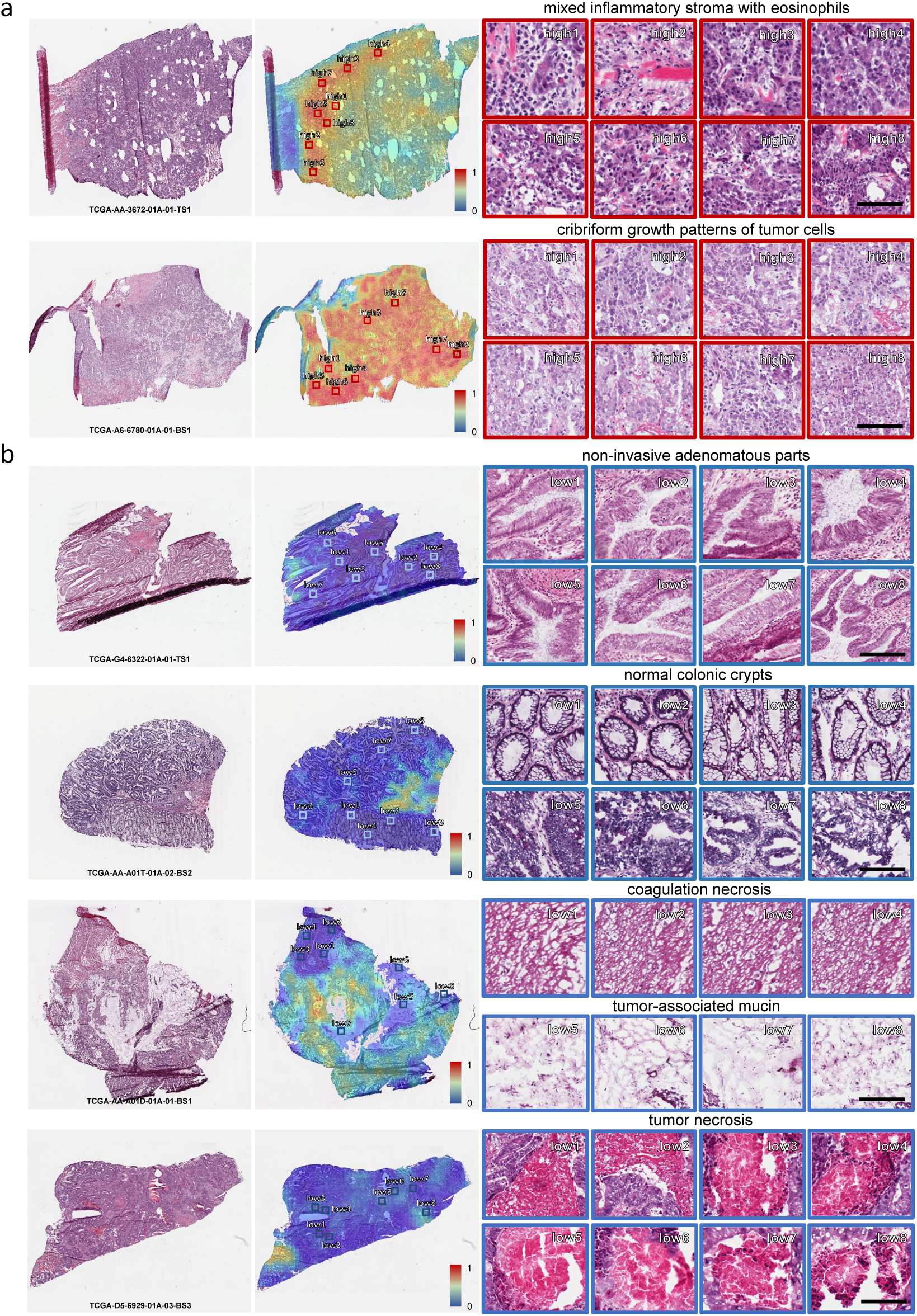
Typical patterns for PDL1 high/low expression in H&E slide images of COAD. **a**, Example slides with typical PDL1 positive and **b,** negative patterns. From left to right are the original H&E slides, predicted heatmaps and 43 example tiles with high and low predicted PDL1 expression. Tiles of high expression are marked in the color red and low ones in blue. Scale bars: 100 *µ*m.

Further, the predicted patterns observed of PDL1 were also validated using paired IHC slides. Correlation analysis between model predictions and IHC scores was conducted using a set of 20 colon adenocarcinoma samples. Visually, PDL1 IHC levels exhibited similar patterns PDL1 as H&E based predictions (Fig. 5a). IHC levels were quantified in patches of 128*×*128 µm^2^ and compared to the H&E based PDL1 prediction in matching areas containing 1% of the total patches on the slide. Model predictions exhibit a consistent strong positive correlation with IHC across all 20 slides, with an average Pearson’s correlation coefficient of 0.74 (Fig. 5b). Together these findings confirm the model’s ability to deconvolve gene expression signals and attribute these signals to distinct histopathological areas. More details are provided in the supplementary material.

**Fig. 5.**
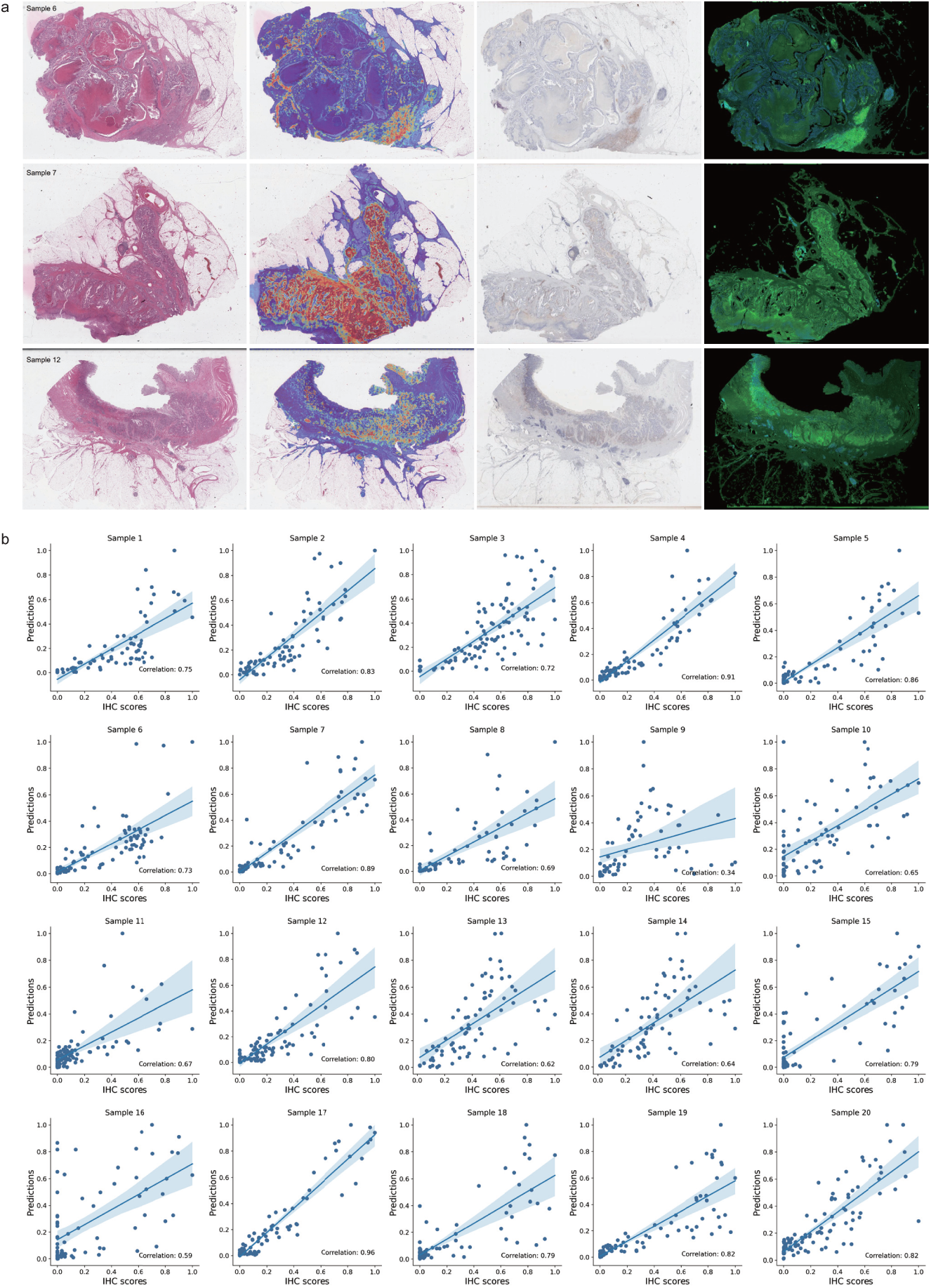
Correlation analysis between model predictions and paired IHC quantification. **a**, Visual comparison between predicted heatmaps and corresponding IHC slide images. The stained-separated images are produced by employing the diaminobenzidine and hematoxylin channels from IHC slide images as the green and blue components, respectively. A more pronounced green area signifies higher PDL1 expression. **b,** Scatter plots illustrating the relationship between normalized IHC quantification and predicted positive probability by the proposed model. The error band represents a 95% confidence interval, calc4u4lated using bootstrap methods. Source data are provided as a Source Data file.

### Correlations between MILTS predictions with TME and clinical features

In order to better understand the observed histopathological associations, correlations with the immune microenvironment, as estimated by CIBERSORT [52] from RNA-seq data, and other clinical parameters were performed. The analysis of immune infiltrates revealed an overall positive correlation between the presence of cell types, such as M1 macrophages and CD8+ cytotoxic T cells, and predicted PDL1 expression across most cancer types (Fig. 6a). This finding aligns with the observed patterns of high PDL1 expression in COAD which was found to co-occur with a mixed inflammatory infiltrate. The analysis also revealed a correlation between the presence of CD4 memory-activated T cells and elevated PDL1 expression, which may indicate a previous immune response against tumor antigens, leading to the upregulation of PDL1 as a countermeasure by tumor cells to suppress T cell activity and evade immune attack. This correlation may imply an intricate interplay between PDL1 expression, immune cell infiltration, and the inflammatory response within the tumor microenvironment.

**Fig. 6.**
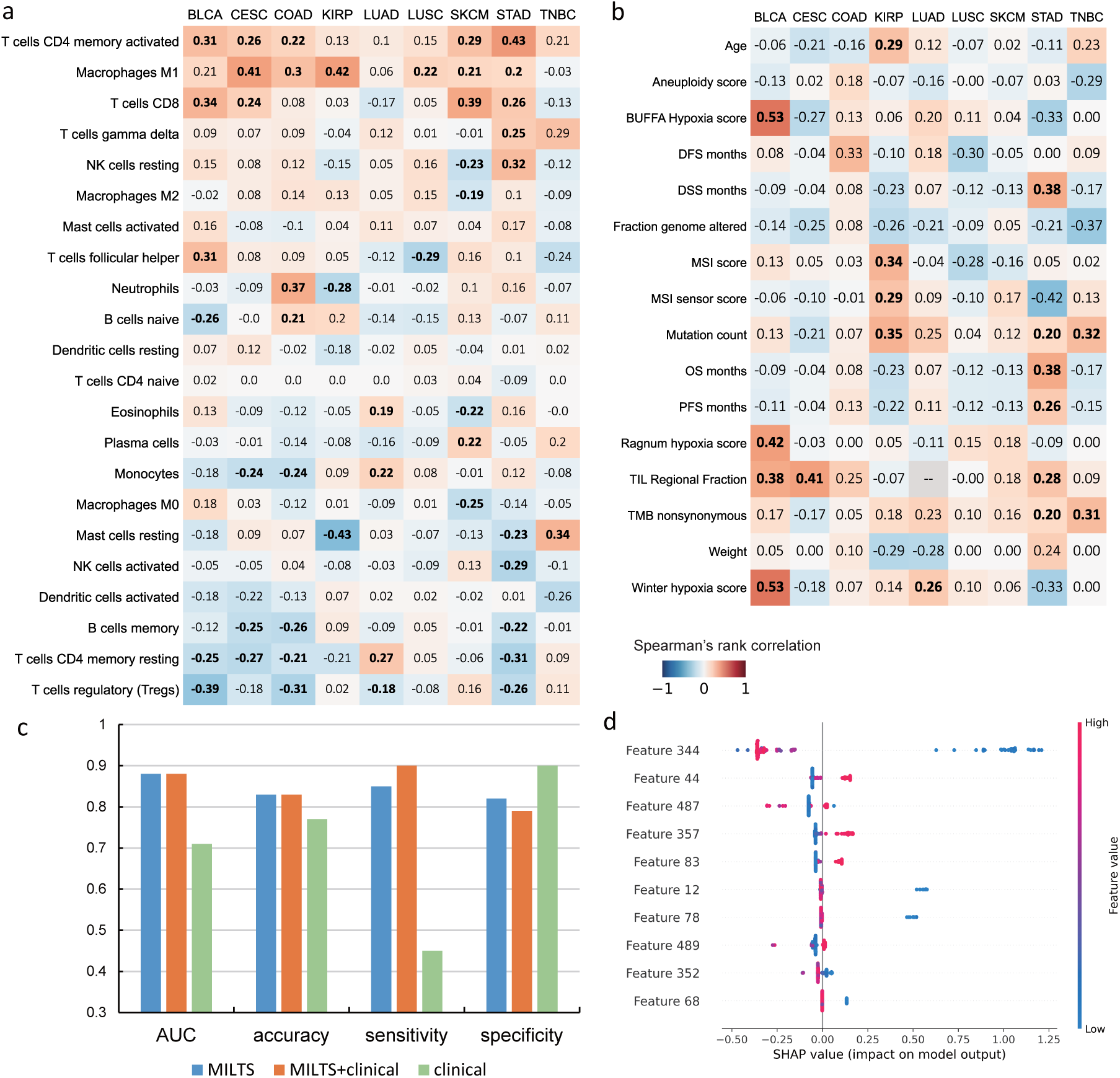
Correlation analysis on tumor immune microenvironment and clinical profiles. **a**, Barplots of Spearman’s rank correlation between model predictions and immune infiltrates. **b,** Heatmap showing Spearman’s rank correlation between model predictions and various clinical features. The bold values indicate correlation coefficients with p-values less than 0.05. The alternative hypothesis is specified as two-sided. **c,** Classification performance by the deep features, deep features concatenated with standardized clinical features and clinical features alone. **d,** Features with the top 10 SHAP values. Features with indices less than or equal to 523 represent deep features extracted by MILTS. Source data are provided as a Source Data file.

In addition, we conducted an analysis to assess the correlation between the predictions generated by MILTS and various clinical features specific to different cancer types which is shown in Fig. 6b. Across cancer types, only few consistent trends were observed, with tumor mutation burden and TIL Regional Fraction exhibiting generally weak positive correlations with PDL1 prediction patterns. The finding of PDL1 expression coinciding with high inflammation, which is often found in tumors with high mutation burden, agrees with the observations of the previous section. To further verify the correlation, we crafted three distinct feature sets, consisting of (i) the deep features derived from the proposed model, (ii) deep features concatenated with standardized clinical features and (iii) clinical features alone. An XGBoost classifier was employed to conduct the prediction based on the forementioned embeddings. The results are presented in Fig. 6c. It’s evident that the deep features extracted by MILTS outperform the purely clinical features which implies that the proposed model encapsulates unique morphological information sourced directly from the pathological slides. We also implemented SHAP to showcase the relative importance of the deep features in comparison to clinical features (Fig 6d). Using SHAP values, it could be observed that the deep features derived from the proposed model hold significantly greater importance than the clinical features. These evidences indicate that the features extracted by the model are mostly either independent of, or lowly correlated with, standard clinical features. Consequently, it suggests that the features captured by the model are distinct and do not overlap with or duplicate the information provided by the clinical features. This highlights the complementary nature of the model’s predictive capabilities in relation to the clinical characteristics of the cancers under investigation.

### Weaker histopathological associations in other cancer types

In order to provide broader context, we also conducted analysis of tumor types, for which immunotherapies have not been approved with PDL1 companion tests, or for which the prognostic significance of PDL1 expression levels has not been significantly verified. The experiments involved a total of 11 types of tumors, which were respectively adrenocortical carcinoma (ACC), esophageal carcinoma (ESCA), head&neck squamous cell carcinoma (HNSC), liver hepatocellular carcinoma (LIHC), mesothelioma (MESO), ovarian serous cystadenocarcinoma (OV), prostate adenocarcinoma (PRAD), rectum adenocarcinoma (READ), testicular germ cell tumors (TCGT), thyroid carcinoma (THCA) and uterine corpus endometrial carcinoma (UCEC). The statistical performance of MILTS on these tumors is presented in Fig. 7a. The average AUC value is measured at 0.67 and the average accuracy stands at 64%, indicating a moderate level of discriminatory power in distinguishing between different mRNA expression levels. See Supplementary Tables S8-S10 for the details.

Among the 11 cancer entities analyzed, it is important to note that not all tumors display a distinct morphological pattern of PDL1 expression that can be directly comparable to PDL1 relevant tumors. The AUCs for HNSC, READ, TGCT and THCA exceeded 0.7, which were comparable to the performance on cancers with PDL1 as an established biomarker where TNBC is the only one with the AUC lower than 0.7. Besides, we also devised a similarity measure based on the distribution of average classification performance of all PDL1 relevant tumors (see Similarity Measure in the supplementary material). The corresponding results are exhibited in Fig. S11, and the four aforementioned tumor types (HNSC, READ, TGCT, and THCA) still demonstrate good similarity. This implies that the intermediate processes connecting macroscopic changes in tumor morphology to microscopic gene expressions may share certain common features for these tumors with good predictability of PDL1 expression, which contribute to their consistent and accurate prediction of PDL1 expression.

**Fig. 7.**
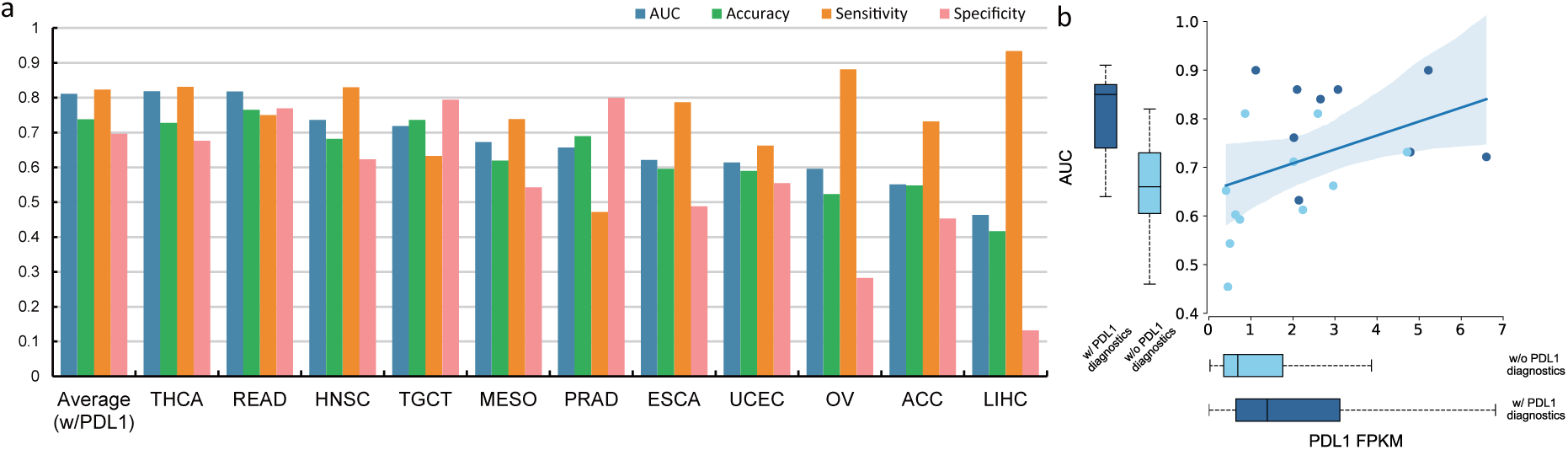
Performance on other cancer entities. **a**, The histogram of model performance on the other 11 cancer entities indicated by AUC, accuracy, sensitivity and specificity. The average performance of the group with PDL1 diagnostics is displayed in the leftmost column. w/PDL1 means the tumors with PDL1 as an established biomarker. **b,** The scatter plot showing the correlation between the overall PDL1 expression level and AUC performance. Box plots displayed alongside the Y and X axes, represent the distribution of tumor-specific AUCs and the PDL1 FPKM, respectively. The central line in each box represents the median value. The box spans the interquartile range (IQR), with its lower and upper boundaries marking the 25th and 75th percentiles, respectively. Whiskers represent the maximum and minimum values. In the AUC box plots, the groups are divided into those with PDL1 diagnosis (*n* = 9) and those without (*n* = 11). For the PDL1 FPKM box plots, the sample sizes are *n* = 3830 and *n* = 3386 for aforementioned two groups. Groups with and without PDL1 diagnosis are distinguished by dark blue and light blue markers in the data. w/, with; w/o, without. The error band represents a 95% confidence interval, calculated using bootstrap methods. Source data are provided as a Source Data file.

The overall distribution of PDL1 expression level of these tumors is shown in Fig. 7b along with that of the PDL1 relevant tumors, where the mean values for two groups are respectively 1.71 and 3.27. From the perspective of classification metrics, it is observed that the mean AUC of these tumors demonstrate a weaker correlation between morphology and mRNA expression level compared to the PDL1-relevant tumors, where a subtle positive correlation exists between the average level of PDL1 expression and corresponding classification performance (Fig. 7b). This implies that the distinction in the predictability based on histopathology may be attributed to multiple factors in addition to PDL1 expression level, which may also have implications for therapeutic outcomes. The observed differentiation of these tumors in morpho-transcriptomic links suggests that additional investigation is warranted for tumors with better morphological correlation.

## Discussion

PDL1 has been proven to be an effective biomarker for determining the response to the immunotherapies in a number of cancer types. Here, we developed MILTS, a novel weakly-supervised multiple instance learning algorithm for predicting slide-level labels from H&E slide scans. We demonstrated its efficacy in predicting elevated PDL1 expression for a broad range of cancers including 9 cancers where PDL1 expression served as an established biomarker. Given the absence of a clinically optimal threshold, different cutoffs thresholds of mRNA quantification were employed in this study to verify the correlation. Compared to existing tools MILTS displayed competitive performance. Our analysis revealed that salient links between histopathology and the level of PDL1 expression for these tumors across various thresholds. Of note, tumor types for which PDL1 is not considered to be a relevant biomarker, also exhibited lower histopathological predictability, likely because of overall lower PDL1 expression.

In addition to the predictability of PDL1 based solely on histology, which could potentially help reduce the need for additional tests, this study also highlighted a diversity of histopathological patterns associated with high and low PDL1 expression. Utilizing the deep histopathological features and differentiated patch-level predictions, some typical patterns for high PDL1 expression in colon adenocarcinomas were demonstrated which include a mixed inflammatory stroma characterized by a relatively high abundance of eosinophils and a cribriform growth pattern of tumor cells. Furthermore, low expression patterns include tumor necrosis, coagulation necrosis at the sample margin, tumor-associated mucin in case of mucinous carcinomas, normal colonic crypts adjacent to the invasive carcinomas, and, most interestingly, non-invasive adenomatous parts of the lesion. It reflected that this weakly-supervised learning manner was capable of pinpointing the potential morphological patterns or cytomorphology of different PDL1 expression levels, which due to their diverse nature and heterogeneous occurrence across and within a slide are difficult to establish by conventional means.

Despite these insights gained from this study, there were still certain limitations. The first one was the persisting performance gap between fresh-frozen and FFPE tissue sections in the proposed model. This discrepancy is likely due to the fact that fresh-frozen slides are generally considered to better preserve the structure of molecular content, as noted by Yu et al. [26]. Therefore, future research should focus on bridging this gap and enhancing the generalizability of models trained with single-modality data. This improvement is crucial for making such models applicable and effective in routine clinical settings. Another important limitation was the consistency of the quantification scale used for supervised information. The accuracy and reliability of the results heavily depended on the quantification method employed to generate supervised information, such as mRNA expression levels in this study. The potential mismatch between the bulk RNA sequences and the tissue presented in the histopathology slide can introduce challenges in integrating the molecular information from bulk RNA sequencing with the spatial information provided by histopathology slides. And variations in the protocols and techniques used for RNA expression quantification can also impact the knowledge transferability between different datasets. And it was believed that these are plausible factors that could contribute to the observed performance differences in the external validation on the CPTAC dataset in this study.

Overall, our study revealed that it is possible to predict high and low PDL1 mRNA expression based on H&E slides with the proposed weakly supervised method MILTS. Furthermore, MILTS captures and identifies meaningful histopathological patterns that are associated with molecular changes, which may not be readily discernible due to their heterogeneous occurrence across and within slides. These findings underscore the utility of AI to augment the capabilities of pathologists by leveraging large amounts of digitized histopathological slides with slide-level annotations, and thereby providing valuable insights into the complex interactions between histology and molecular biology.

## Methods

### Datasets

Whole slide images and transcriptome profiling data utilized in the experiments comes from TCGA project via National Cancer Institute (NCI) Genomic Data Commons Portal [53] and CPTAC project via the Cancer Imaging Archive (TCIA) Pathology Portal, which comprises 12,299 H&E stained whole slide images and corresponding mRNA quantification data obtained from 6,715 patients diagnosed across 20 different types of cancer. 20 paired FFPE and IHC colorectal cancer samples are obtained from Charité–Universitätsmedizin Berlin. Data from CPTAC dataset is employed for external validation purposes. The sample types comprise mainly primary solid tumor as well as metastatic tissues. Specifically, the dataset comprises 8,719 fresh-frozen slides and 2,966 FFPE slides from TCGA, along with 614 slides from CPTAC COAD and BRCA. Characteristics of patients is presented in Supplementary Table S11. We collect all available whole slide images scanned at a magnification of 20*×* or higher, along with their CD274 mRNA expression level read counts normalized by the upper quartile fragments per kilobase of transcript per million mapped reads (FPKM-UQ).

According to the biospecimen information available for TCGA cases, it is confirmed that the digitized fresh-frozen slides and sequencing data usually originate from the same vial of the same sample, with the slides typically obtained from either the top or bottom layer of the corresponding section. However, FFPE slides are sampled from a different vial and the potential mismatch with the sequencing data could be more pronounced. Thus a thresholding strategy was implemented on sequencing data converting the task into a classification task rather than a regression one, thereby enhancing the alignment between the data and labels. A commonly used split for data, in which 60% is reserved for training, 15% for validation, and 25% for testing, was employed for the majority of the tumors analyzed. It’s worth noting that the splitting was performed based on patient IDs. This led to a distribution of 5,373 slides for training, 1,189 slides for validation, and 3,182 slides for testing in the case of fresh-frozen slides, and 1,837, 386, and 743 slides respectively for FFPE slides.

### Data preprocessing

Due to the tremendously large size, whole slide images should be decomposed into smaller elements so that deep learning techniques like convolutional neural network could handle them at an acceptable computational cost. Hysteresis thresholding is applied to exclude the invalid background by finding contours of stained tissues. The valid regions within the contours or partly overlapped are cropped into tiles of 256 pixels*×*256 pixels at resolution of 20*×*, which correspond to the physical size of 128 *µ*m*×*128 *µ*m respectively. In total, 52,587,256 tiles are extracted for the establishment and verification of the model. To mitigate the bias introduced by variations in staining protocols and laboratory conditions, we applied random adjustments to the brightness, contrast, saturation, and hue of generated tiles. In addition, random rotation and crop of the tiles were also applied to increase the variability of scales and orientations. Detailed parameter settings for data augmentation are shown in Supplementary Table S12.

### Teacher-student collaborated multiple instance learning (MILTS) for patch feature extraction

A two-stage strategy is adopted in the diagnosis of a patient’s slide, starting with patch-level assessment, followed by slide-level feature aggregation and prediction. This framework enables not only higher-level diagnosis of the patient, but also interpretation of the slide’s inner structure based on the patch-level predictions. Specifically, prediction of PDL1 expression is formulated as a classification problem using a specific mRNA quantification value as the threshold. This threshold was set based on the median point, upper tertile, and upper quartile of the mRNA quantification distribution. Classification based on the patch-level classifier is usually devised under multiple instance learning framework when only the slide-level property is known and potential heterogeneity is implied. During the patch-level feature extraction stage, MILTS made two important improvements. Firstly, we modified the optimization of multiple instance learning framework to a class-targeted way considering the heterogeneous composition of a slide image. Secondly, inspired by the human-decision making process, the single model of MIL is decomposed into a teacher and a student model, in which the teacher model dynamically assigns probabilities to tiles, and the student model subsequently learns to predict. These two components work collaboratively to boost the performance on patch-level pathological feature extraction.

In the classic MIL approaches, the target label *Y_k_* is predicted based on the distribution of the instances (or patches) *{x*_1_*_,k_, x*_2_*_,k_, …, x_n,k_}* from a bag (a slide image in this case) *X_k_*, where the hypothetical distribution should follow the principle below [54]:

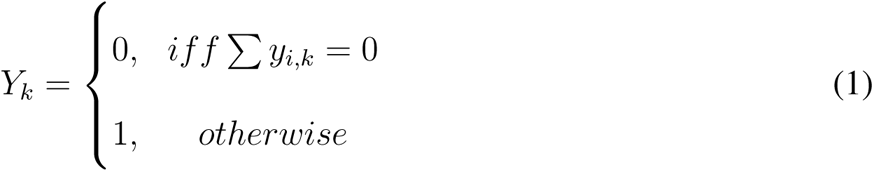

where *y_i,k_*is the label for instance *x_i,k_*and the principle applies to the binary classification. This assumption implies that positive instances should only be present in positive bags, which accurately reflects certain scenarios in pathological diagnosis, such as distinguishing between benign and malignant lesions.

However, it is crucial to note that PDL1 expression is quantified by the accumulation of cancer cell surface PD-L1, rather than simply by its presence or absence. This implies that elements defined as highly or lowly expressed could exist in cohorts with contrary macro statistical properties. Based on this observation, we propose our class-targeted optimization way, which regularizes the instance-level learning by filtrating the representative instances for slides with varying labels. Instances from different slides are selected based on class-representative criteria defined by their respective slide labels. Specifically, a maximum-minimum selection criteria is established to determine the groups of instances involved in model training for binary classification. In the scenario where patients should be classified as high or low PDL1 expression, instances from positive samples with maximum response and negative samples with minimum response tend to be selected and share the same label with bags they belong to. The criteria could be summarized as:

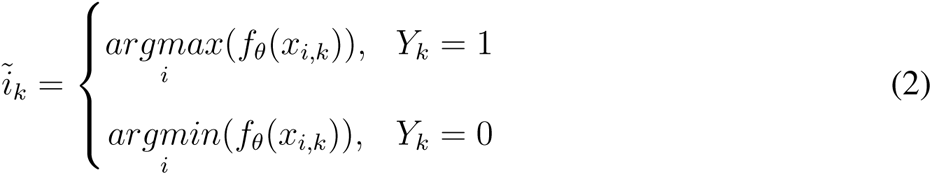

where 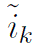 is the index of the selected instance and *f_θ_* is the function of instance-level classifier which maps the input patch *x_i,k_* into a normalized hidden variable indicative of its typicality for the slide. To incorporate additional histological pattern references, the criteria can be further relaxed by allowing *M >* 1 representative instances in the slide.

Building on our improved class-representative MIL approach, we further consider data composition in this task as a mixture of labeled and unlabeled instances and introduce the teacher-student collaborative learning into MIL according to the characteristics of whole slide images and weakly annotations. Typically, the representative instances selected from a slide are considered as labeled data and assigned the same label as the slide. Nevertheless, these instances only represent a small portion of the entire slide image. Relying solely on these instances for training can result in overfitting and suboptimal performance due to their limited representation of the entire slide image. There still exist a considerable number of instances within whole slide images that also contain pathological information and can be utilized for representation learning. Usually to address the challenge of interpreting slides containing both clear and ambiguous regions, a pathologist typically relies on their own expertise to interpret the clear parts of the slide (labeled) and consults with more experienced or senior pathologists for guidance on the ambiguous regions (unlabeled). And this process also benefits senior pathologists by exposing them to difficult cases. Taking inspiration from this process, we propose to decompose the single model in the MIL framework into a temporal ensemble model consisting of a teacher model, acting as the senior pathologist retrieving regularization of unlabeled data on feature representation, and a student model, learning from the teacher model. The mean teacher method [55] is adopted to constructed the teacher model and ResNet34 [56] as baseline architecture for teacher&student model. Specifically, labeled instances *X_labeled_* = *{x*_1_,_1_*, x*_1,2_*, …, x_K,_*_1_*, …, x_K,n_}* selected by the teacher model, where *K* instances of each slide are assigned with the label consistent with the sample, and unlabeled instances *X_unlabeled_*= ∁*_X_X_labeled_* are both inputted into student model and compute a probability of high PDL1 expression in every iteration of training. The weighted cross-entropy loss function, which is used as the classification cost, penalizes the difference between the output and pseudo label for the clear regions of the slide represented by *X_labeled_*. Meanwhile, the consistency cost is applied to the ambiguous regions represented by *X_unlabeled_* and constrains the distance to the distribution given by the teacher model. The loss function was formulated as:

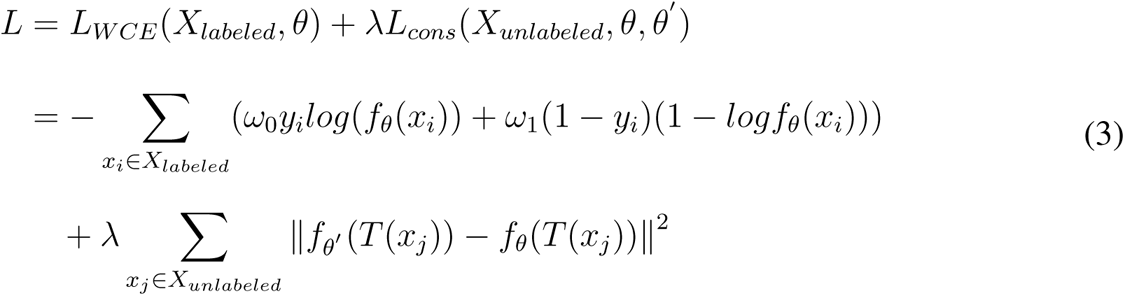

where *ω_i_*is the weight to balance the class frequency and 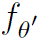 is the function of the teacher model generated by exponential moving average (EMA) of previous student models. *T* () is a combination of transforms including random rotation and color jitter. The parameters of the teacher model would be updated after the student model is optimized in each iteration.

In other words, the proposed approach constructs a teacher model to assess the reliability of knowledge and pairing the reliable parts with corresponding answers for the student model to learn. For data with greater uncertainty judged by the teacher model, the student model will attempt to imitate the teacher model’s response even if an exact answer is not available. The teacher model will then use feedback from the student model to update itself and improve its own knowledge. Among different epochs of training, labels are dynamically assigned for instances according to the predicted probability of the teacher model via the above mentioned MIL manner, where meaningful histopathological patterns are expected to be uncovered through aggregated information distilled from mean teacher. And the student model optimized by gradient descent with regard to objective function *L* would promote the prediction of the teacher model in turn by a weighted average behavior over training steps. This iterative process allows for continuous learning and refinement of both the teacher and student models to capture the effective histopathological features associated with PDL1 expression. Usually both models could be utilized to output the positive probability at the end of the training which share similar performance for patch-level prediction.

### Enhancing patient-level diagnosis by fused slide tokens with statistical summary of patch-level predictions

In the slide-level feature aggregation stage, statistical summary of patch-level predictions are merged with slide tokens by leveraging the local details extracted by the patch-level CNN and the global perspective of a transformer model. Given the trained model 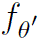 or *f_θ_* at patch level, the typicality of all effective patches in a slide would be computed, which could be further interpreted as positive or negative probability according to the slide-level property. Based on the patch-level prediction, several statistical features of the slide were extracted including percentage of positive patches, histogram of probability distribution, median value and mean value of positive probability. Specifically, patches with typicality over 0.8 and below 0.2 are removed when computing the above features because a trimmed estimator is supposed to better reflect the central tendency of data. Further, features *e* = [**e_1_**, **e_2_***, …,* **e_m_**] extracted by the patch-level backbone (i.e. feature vectors output by the adaptive pooling layer of ResNet34 in the end) are aggregated to form the global representation of the slide through the attention mechanism. Transformer is utilized to fuse these features into a class token 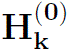 which could be formulated as [46]:

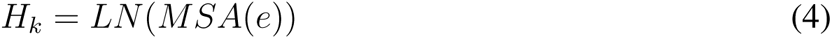

where *LN* (*·*) is layer normalization and *MSA*(*·*) indicates multi-head self attention operation. The first row vector of correlation matrix 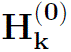 i.e. class token is taken as the aggregation of all local features in a slide. Training details follow the settings in [46]. Along with the statistical features mentioned above, the representation of the slide **E** *∈* R^1^*^×^*^523^ was constructed by concatenation and conducted the diagnosis by using MLP to classify the slide representation as PDL1 high expression or low expression. The MLP model is trained with cross-entropy loss and SGD optimizer, where the learning rate is set as 2 *×* 10*^−^*^4^.

### Implementation details

Data preprocessing mainly including segmentation and patching of whole slide images was implemented on multiple workstations considering massive amount of data, whose central processing units are respectively AMD Ryzen 5950X, Intel i9-12900K and Intel i9-10900K. Our weakly supervised framework MILTS and patient-level feature aggregation&diagnosis models were then trained and evaluated on a platform of GIGABYTE GeForce RTX 3090Ti with 24 GB of graphics memory. Algorithms were mostly programmed with Python (version 3.8.10) and libraries mainly involved were OpenCV (version 4.5.3), OpenSlide (version 1.1.2), Pillow (version 8.4.0), scikit-image (version 0.18.3) and Numpy (version 1.19.5). We utilized Pytorch [57] as the basic machine learning framework for data loading, enhancement, training and inference pipeline. Before the start of MILTS training, we initialized ResNet34 with ImageNet [44] pretrained parameters for both teacher and student model. A mini-batch size of 512 was adopted to accelerate computation and we used a stochastic gradient descent (SGD) optimizer with an initial learning rate of 1 *×* 10*^−^*^2^ to optimize weights of the models. The proportions of labeled instances within a single slide were defined as 0.25, 0.35, and 0.45 for the quartile, tertile, and median points, respectively. The consistency cost weight *λ* for the loss function was determined to be 100. EMA decay *α* for updating the teacher model was set to 0.99 throughout training. The strategy of cosine annealing was applied to schedule the learning rate and the minimum value was set as 1 *×* 10*^−^*^4^. All the models were trained for 30 epochs, during which two circles of learning rate change were completed. On our platform using a single GPU, the typical inference time for a 20*×* whole slide image with non-overlap patches was 6.96s, which indicated a high efficiency and may be further promoted by large-scale parallel computing.

## Data availability

The TCGA data (images, as well as transcriptomic and clinical data) used in this study are publically available from http://gdc.cancer.gov. CPTAC image data are publically available from https://wiki.cancerimagingarchive.net/display/Public/CPTAC+Imaging+Proteomics and transcriptomic data from http://gdc.cancer.gov. The imaging data from Charité–Universitätsmedizin Berlin is accessible upon request. Source data are provided with this paper.

## Code availability

The code in this paper are available through a Code Ocean compute capsule (https://codeocean.com/capsule/2580510/tree).

## Acknowledgments

This work was supported by grants from the National Natural Science Foundation of China under Grant 62271016, in part by the Beijing Natural Science Foundation under Grant 4222007, in part by the Fundamental Research Funds for the Central Universities and in part by the scholarship programme of the China Scholarship Council under Grant 202206020151. The laboratory of T. G. P. G. is supported by the Barbara and Wilfried Mohr Foundation. We would also like to extend our thanks to Areeba Patel, Yiheng Tang, Heng Luo, Domenico Calafato, Gleb Rukhovich, Artem Lomakin and Anna Mathioudaki for helpful discussions and assistance in data processing.

## Author Contributions Statement

D. J., M. G. and X. B. conceived the study. D. J. developed the deep learning algorithms. D. J., S. L., A. S. and T. G. P. G. performed the experiment analysis. T. G. P. G., D. H. and A. A. prepared and collected pathology slides. T. G. P. G. and D. J. conducted the histopathological analysis. D. J., M. G. and X. B. wrote the manuscript with help of all other authors. M. G. and X. B. supervised the project.

## Competing Interests Statement

The authors declare no competing interests.

## Notes

### Competing Interest Statement

The authors have declared no competing interest.

### Summary of Updates

Author affiliations updated; Abstract revised; Results updated with supplemented experiments results on FFPE slides and external validation; Figure 1, 4, 6 revised; Figure 5 added;

