## supplementary figures and tables for "Teacher-student collaborated multiple instance learning for pan-cancer PDL1 expression prediction from histopathology slides"

**Correlation analysis between model predictions and paired IHC slides.**

To conduct the validation on IHC imaging, we collected 20 colon adenocarcinoma samples with paired H&E slides and IHC slides. In particular, we employed a color deconvolution technique [1] to separate the immunohistochemical staining from the hematoxylin counterstaining, allowing us to quantify PDL1 expression in the IHC slide images as shown in Fig. S1. The diaminobenzidine (DAB) channel was extracted from the original IHC tiles, and its pixel-wise intensity was utilized for the quantification of PDL1 expression. Subsequently, we performed image registration between the H&E and IHC slide pairs using SIFT [2] as the feature descriptor to mitigate any shifts and distortions (Fig. S2).

With the registered image pairs in place, we proceeded with regional-level correlation analysis. The entire slide images were partitioned into 100 regions, and Pearson's correlation coefficient was calculated by accumulating the predicted positive probability and diaminobenzidine intensity within each region. The predictions by the model trained at the tertile threshold with COAD FFPE data were adopted here.

**Similarity measure between PDL1 relevant tumors and other tumors.**

To explore the similarity between the aforementioned tumors and the PDL1-relevant ones, chi-square distribution was utilized to model the distribution of classification metrics values for PDL1-relevant tumors. A bootstrapping strategy with 100 random resamples was applied to each of the 9 tumors thus 900 subsets were constructed, whose classification metric values were utilized to construct the template matrix. Mahalanobis distance is used to measure the similarity between the cluster and each single point from the cluster. In this manner, the average distances between PDL1 relevant tumors and the template matrix at the threshold of median, upper tertile and upper quartile are respectively 1.849, 1.769 and 1.747. A chi-square distribution well fitted the aforementioned distance distribution, whose parameters could be found in supplementary Table S13. Corresponding P values through two-sided Kolmogorov-Smirnov (KS) test are 0.581, 0.172 and 0.190. The specific distributions are presented in supplementary Fig. S10. As the distance is regarded as continuous random variable, it is more appropriate to calculate corresponding probability density rather than the probability for individual data points. The probability density values listed in descending order are respectively 0.502 for HNSC, 0.481 for MESO, 0.450 for TGCT, 0.358 for UCS, 0.350 for THCA, 0.326 for READ, 0.280 for ACC, 0.280 for ESCA, 0.278 for OV, 0.258 for PRAD and 0.120 for LIHC as presented in Fig. S9, which could be taken as a similarity measure to the template matrix. Based on the measurement of correlation between morphological patterns and PDL1 mRNA expression levels, it was observed that some tumors such as HNSC, TGCT, THCA and READ still exhibit a high similarity to PDL1-relevant tumors. The observed differentiation in morpho-genomic links suggests that additional investigation is warranted for tumors with better morphological correlation. This implies that the intermediate processes connecting macroscopic changes in tumor morphology to microscopic gene expressions may share certain common features for these tumors and PDL1-relevant ones.

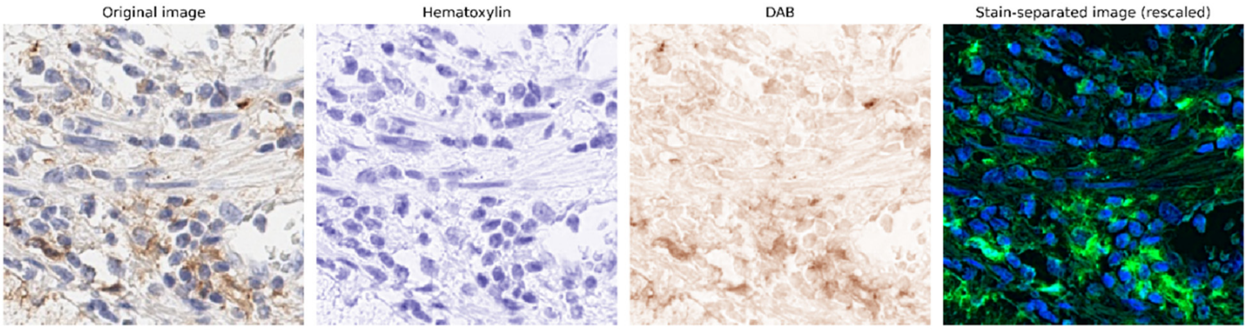


**Fig. S1. The original IHC tile, hematoxylin channel, DAB channel and the pseudo fluorescence image generated by the two abovementioned channels.**

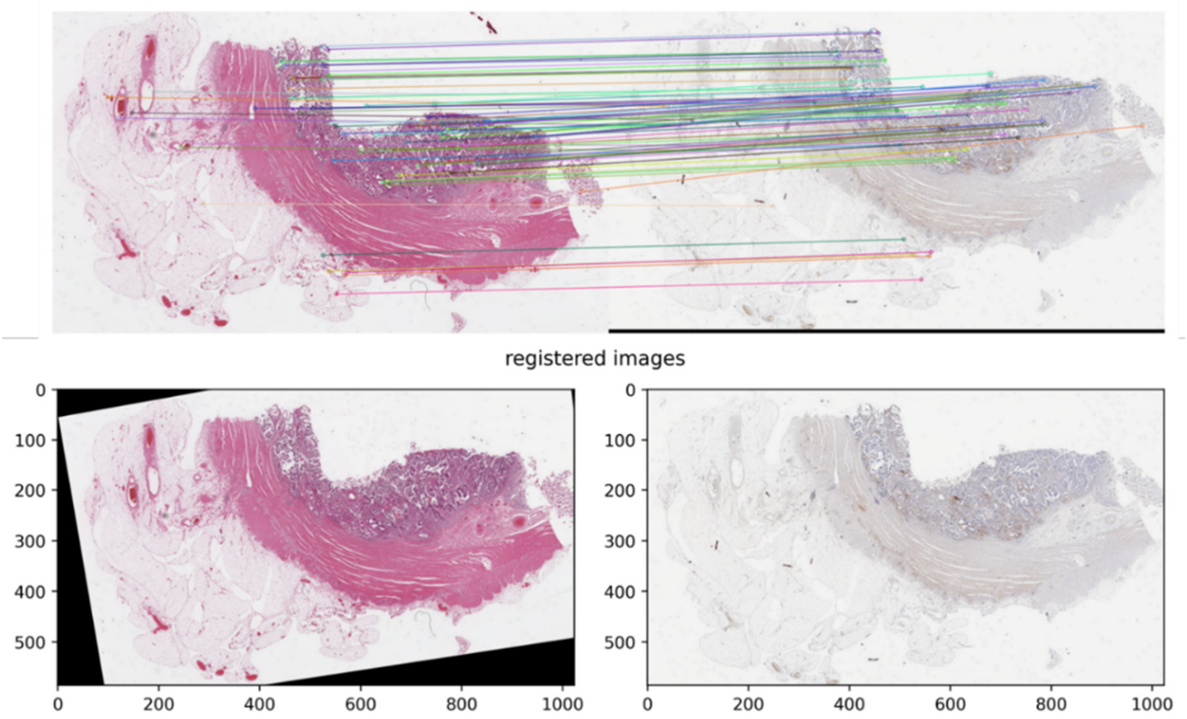


**Fig. S2. Example of registration between the H&E slide image and the corresponding IHC slide image.**

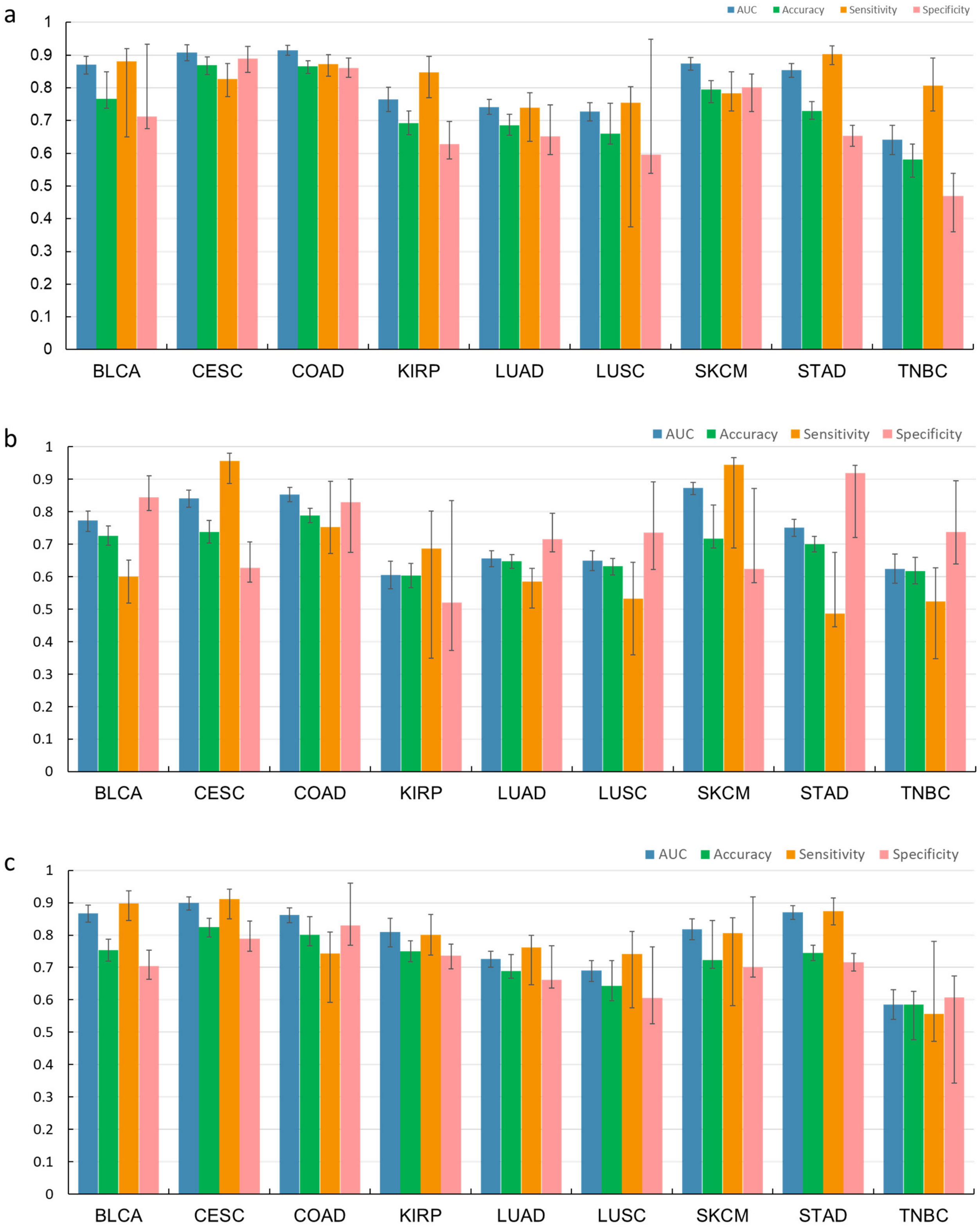


**Fig. S3. Classification performance on PDL1 clinically relevant tumors obtained at different thresholds**. **a**, Results at the threshold of upper tertile. **b**, Results at the threshold of median point. **c**, Results at the threshold of upper quartile. The data partitioning settings at the threshold of upper quarter and median point follow those at the threshold of upper tertile. Error bars represent 95% confidence intervals, calculated using bootstrap methods.

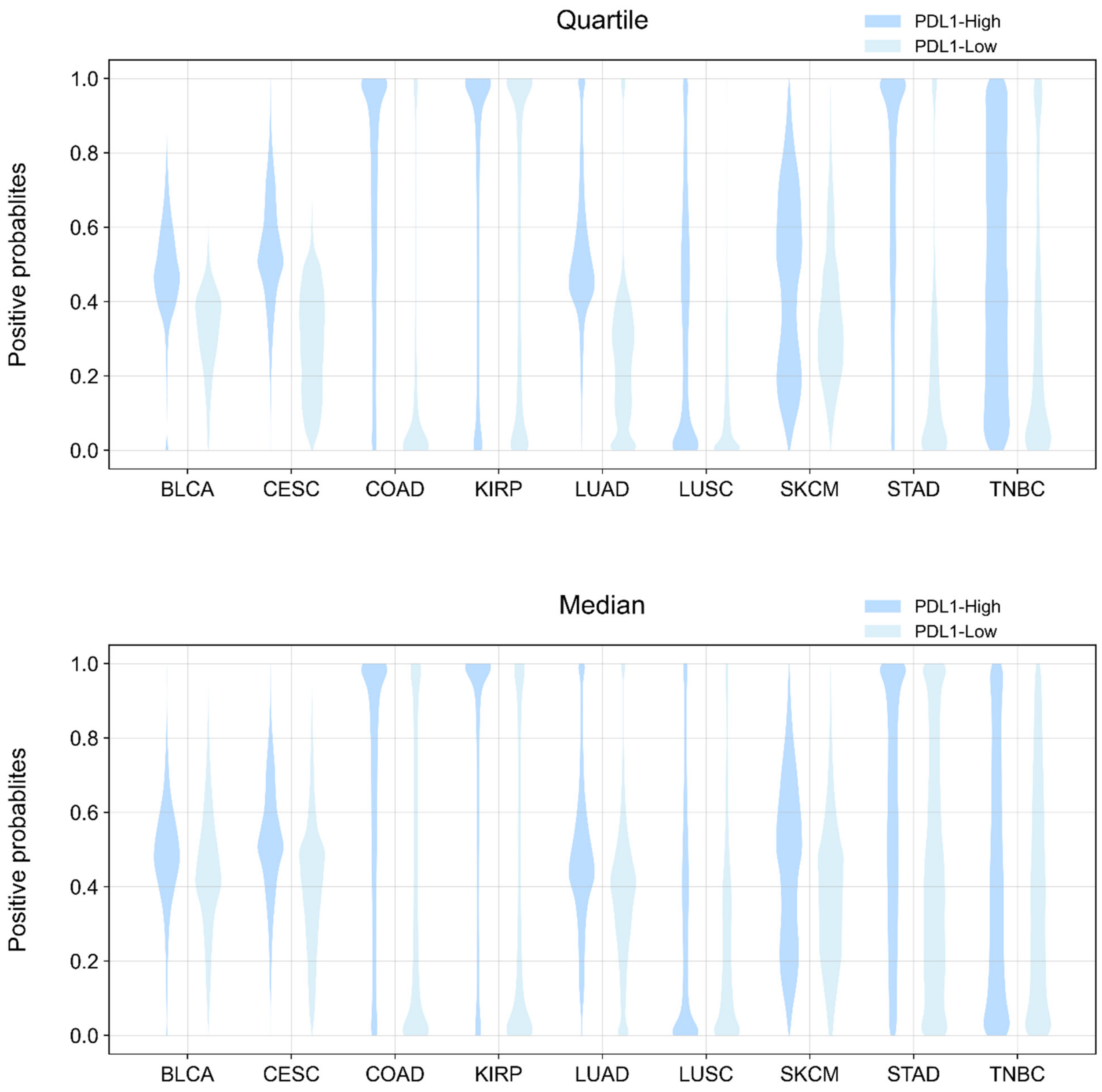


**Fig. S4. Distribution of positive probability predicted by MILTS at the quartile and median thresholds on PDL1-relevant tumors.**

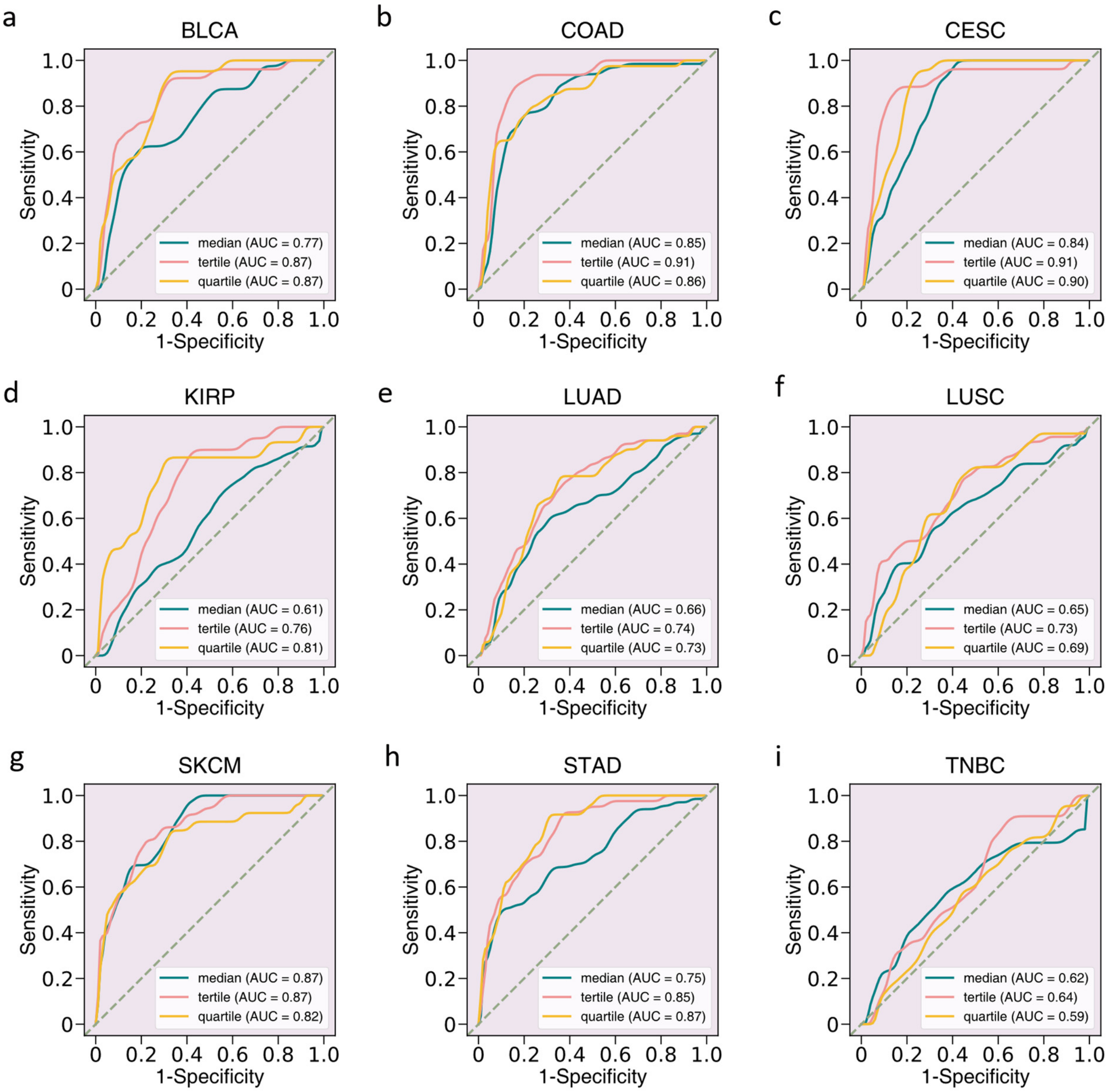


**Fig. S5. Performance and results obtained by MILTS at different thresholds on clinically PDL1-relevant tumors.** The corresponding ROC curves of clinically PDL1-relevant tumors with respect to different thresholds, where green, pink and yellow lines indicate respectively the performance achieved at the threshold of median, upper tertile and upper quartile. The results shown here only involve data of the test set.

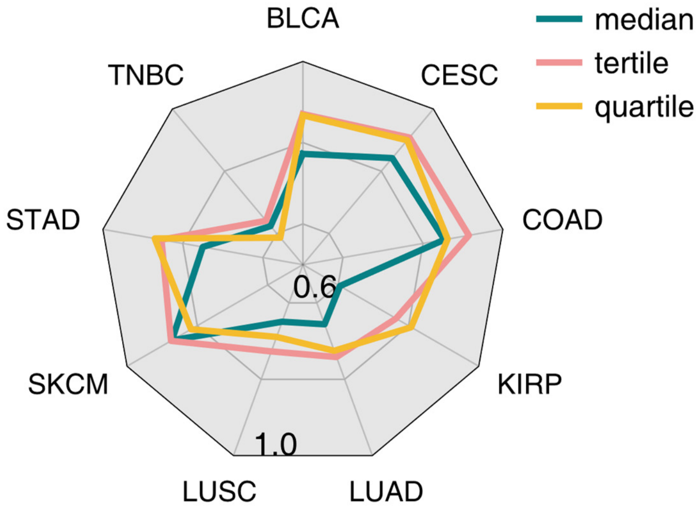


**Fig. S6. AUC obtained by MILTS at different thresholds on clinically PDL1-relevant tumors.** The radar chart demonstrating the AUC distribution in terms of different thresholds, where green, pink and yellow lines indicate respectively the performance achieved at the threshold of median, upper tertile and upper quartile. The results shown here only involve data of the test set.

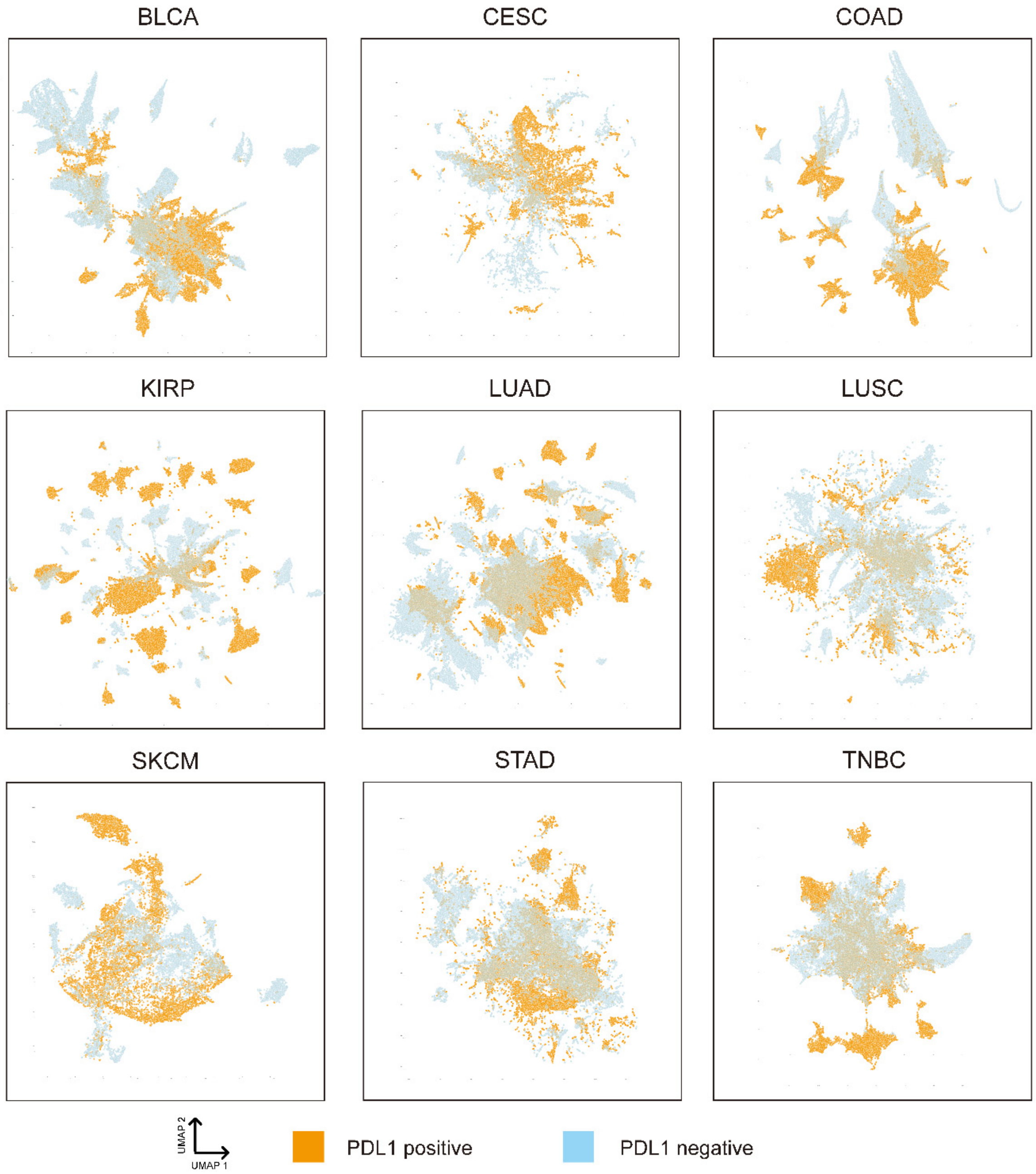


**Fig. S7. Two-dimensional visualization by patch-level embeddings.** Visualization of the representation space constructed by the patch-level classifier of MILTS for test set. The elements in orange indicate the instances from PDL1-high slides while the blue ones are from PDL1-low slides.

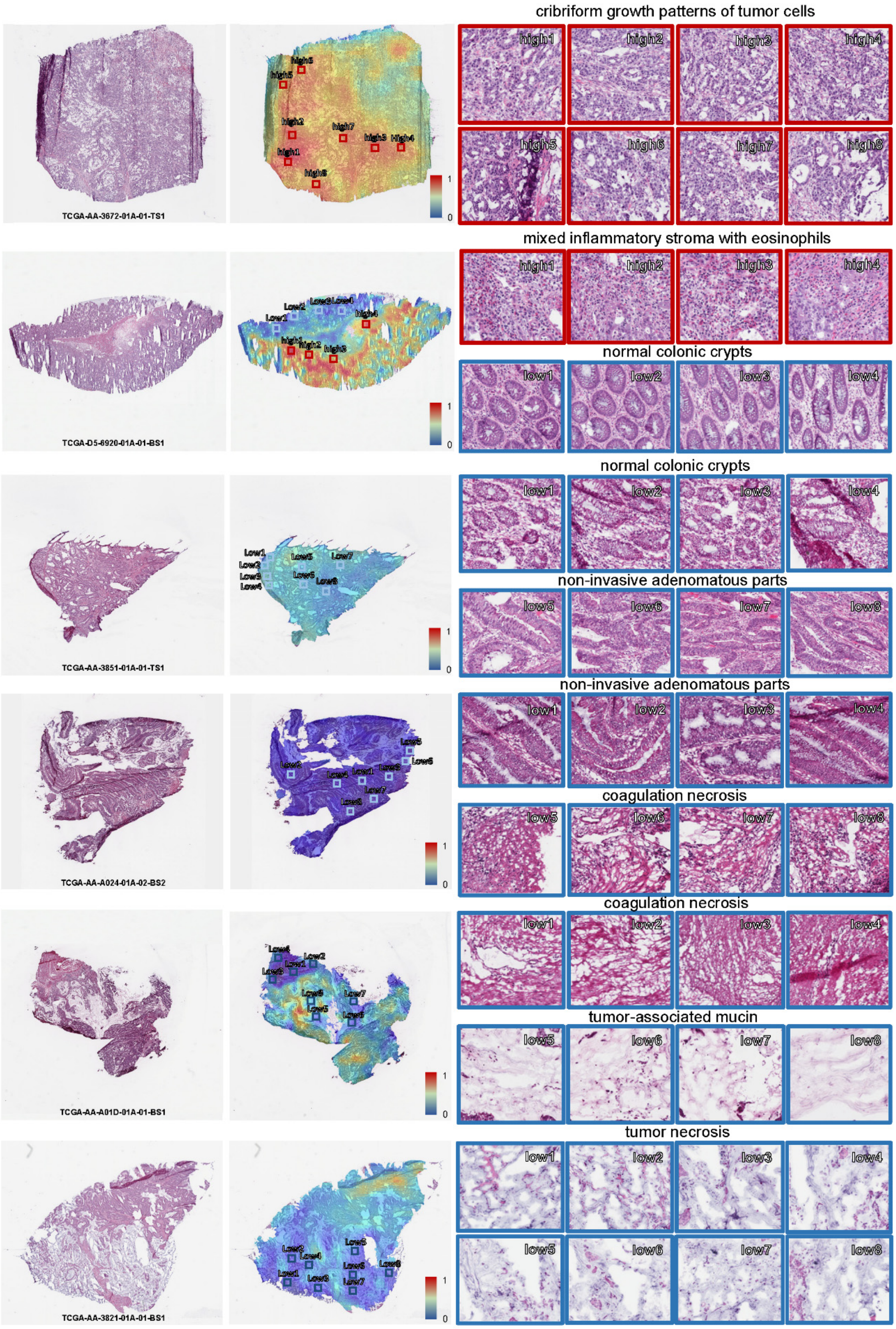


**Fig. S8. Typical patterns for PDL1 high/low expression in fresh-frozen slides of COAD.**

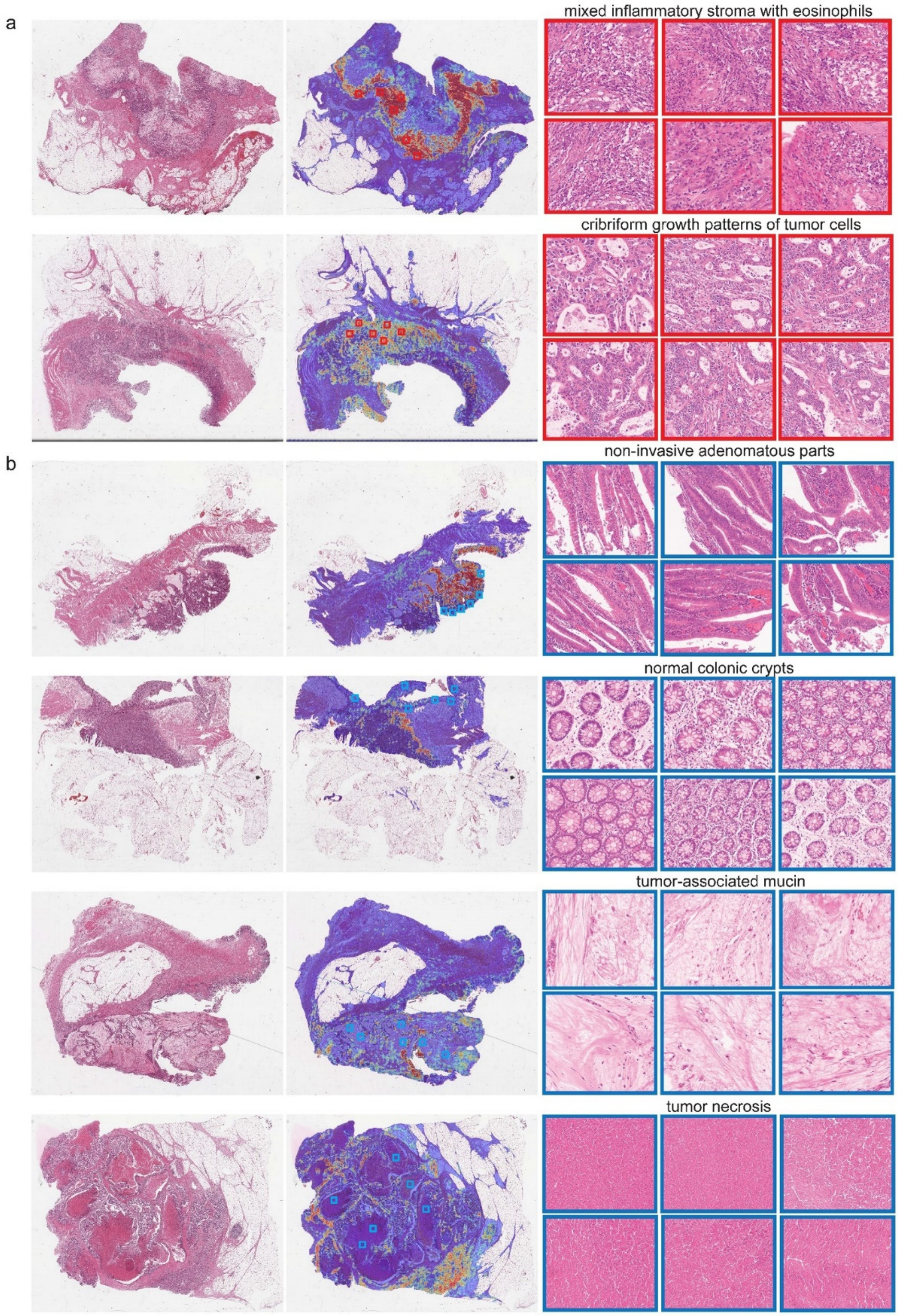


**Fig. S9. Typical patterns for PDL1 high/low expression in FFPE slides of COAD. a,** Example slides with typical PDL1 positive and **b,** negative patterns.

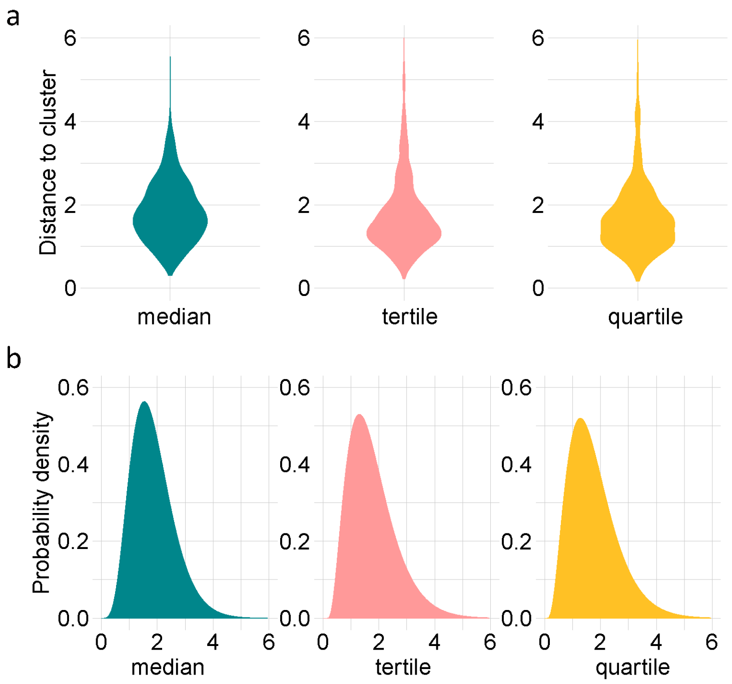


**Fig. S10. The overall distribution of the distance between the PDL1 relevant tumors and the template cluster. a**, The violin plot demonstrating the distribution of the Mahalanobis distance between the template cluster and PDL1-relevant ones at different thresholds. **b**, The probability density curves of the fitted chi-square distribution of the Mahalanobis distance between the template cluster and PDL1-relevant ones at different thresholds.

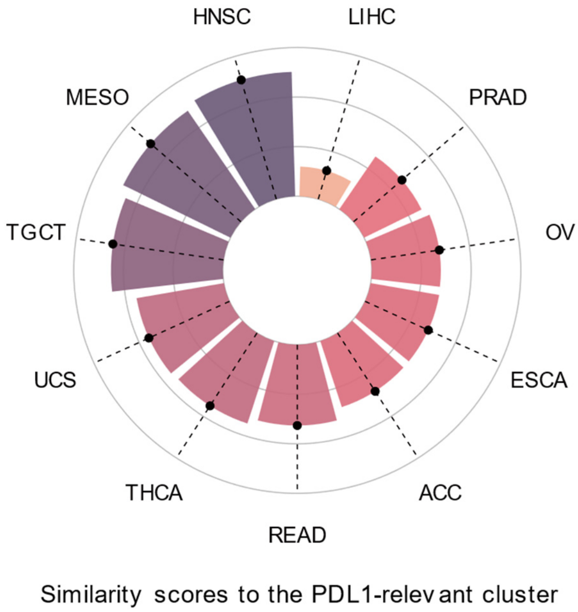


**Fig. S11. Circular bar chart of probability density of tumors being PDL1 clinically-relevant using quantitative results obtained at the mean of three thresholds.**

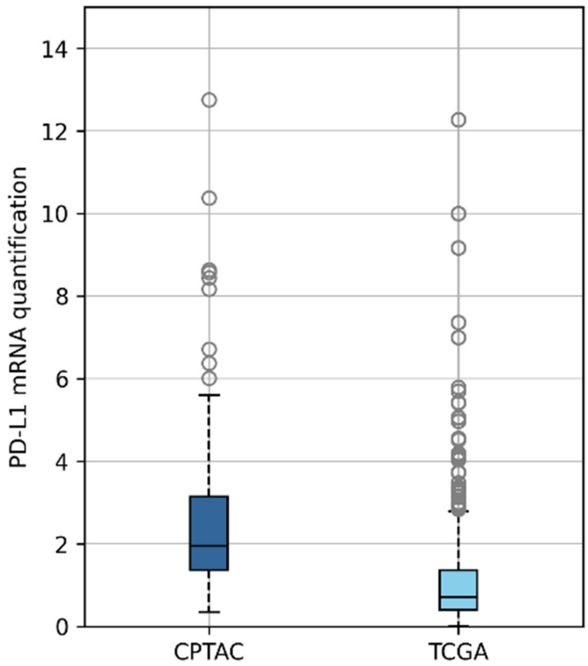


**Fig. S12. mRNA distribution of CPTAC and TCGA COAD dataset.** The central line in each box represents the median value. The box spans the interquartile range (IQR), with its lower and upper boundaries marking the 25th and 75th percentiles, respectively. Whiskers extend 1.5 times the interquartile range from the first and third quartiles. The sample sizes are respectively $n$=100 and $n$=451.

Table S1

QUANTILE VALUES FOR PDL1-RELEVANT TUMORS

| Cancer type | BLCA | CESC | COAD | KIRP | LUAD | LUSC | SKCM | STAD | TNBC |
|---|---|---|---|---|---|---|---|---|---|
| Quartile | 3.0 | 6.1 | 1.4 | 2.2 | 4.7 | 7.2 | 2.3 | 2.2 | 2.3 |
| Tertile | 2.0 | 4.2 | 1.1 | 1.6 | 3.8 | 5.0 | 1.7 | 1.8 | 2.0 |
| Median | 1.0 | 2.4 | 0.7 | 1.2 | 2.5 | 2.9 | 1.0 | 1.2 | 1.4 |

Table S2

QUANTILE VALUES FOR OTHER TUMORS

| Cancer type | THCA | READ | HNSC | TGCT | MESO | PRAD | ESCA | UCEC | OV | ACC | LIHC |
|---|---|---|---|---|---|---|---|---|---|---|---|
| Quartile | 3.6 | 1.1 | 5.0 | 2.6 | 2.4 | 0.8 | 2.6 | 0.8 | 1.1 | 0.5 | 0.6 |
| Tertile | 3.0 | 0.9 | 4.0 | 2.1 | 1.6 | 0.7 | 2.0 | 0.7 | 0.9 | 0.4 | 0.4 |
| Median | 2.2 | 0.6 | 2.4 | 1.3 | 1.0 | 0.6 | 1.4 | 0.6 | 0.6 | 0.3 | 0.3 |

Table S3

QUANTITATIVE RESULTS OF THE PROPOSED MODEL ON PDL1 CLINICALLY RELEVANT TUMORS AT MEDIAN THRESHOLD.

| | LUAD | LUSC | BLCA | TNBC | CESC | STAD | COAD | KIRP | SKCM |
|---|---|---|---|---|---|---|---|---|---|
| AUC | 0.656 (0.631~ 0.680) | 0.650 (0.619~ 0.680) | 0.773 (0.740~ 0.802) | 0.624 (0.580~ 0.669) | 0.841 (0.815~ 0.866) | 0.751 (0.724~ 0.776) | 0.854 (0.831~ 0.876) | 0.605 (0.563~ 0.647) | 0.872 (0.853~ 0.891) |
| Accuracy | 0.648 (0.626~ 0.669) | 0.633 (0.606~ 0.656) | 0.726 (0.696~ 0.756) | 0.617 (0.578~ 0.660) | 0.737 (0.704~ 0.773) | 0.701 (0.676~ 0.723) | 0.789 (0.766~ 0.811) | 0.603 (0.566~ 0.641) | 0.718 (0.689~ 0.821) |
| Sensitivity | 0.585 (0.503~ 0.625) | 0.532 (0.359~ 0.644) | 0.599 (0.518~ 0.650) | 0.523 (0.348~ 0.627) | 0.957 (0.886~ 0.980) | 0.487 (0.445~ 0.676) | 0.753 (0.671~ 0.894) | 0.687 (0.348~ 0.801) | 0.945 (0.688~ 0.967) |
| Specificity | 0.716 (0.676~ 0.795) | 0.736 (0.622~ 0.893) | 0.844 (0.803~ 0.910) | 0.738 (0.639~ 0.895) | 0.628 (0.583~ 0.707) | 0.918 (0.720~ 0.943) | 0.829 (0.674~ 0.900) | 0.521 (0.373~ 0.835) | 0.623 (0.582~ 0.872) |

Data enclosed in the parentheses represent the corresponding lower bound and the upper bound of 95% confidence interval. The intervals were obtained by bootstrapping strategy with 2000 random resamples.

Table S4

QUANTITATIVE RESULTS OF THE PROPOSED MODEL ON PDL1 CLINICALLY RELEVANT TUMORS AT THE THRESHOLD OF UPPER QUARTILE.

| | LUAD | LUSC | BLCA | TNBC | CESC | STAD | COAD | KIRP | SKCM |
|---|---|---|---|---|---|---|---|---|---|
| AUC | 0.725 (0.700~ 0.750) | 0.690 (0.657~ 0.720) | 0.868 (0.841~ 0.892) | 0.586 (0.540~ 0.631) | 0.899 (0.878~ 0.918) | 0.870 (0.848~ 0.890) | 0.862 (0.839~ 0.883) | 0.809 (0.764~ 0.851) | 0.817 (0.785~ 0.850) |
| Accuracy | 0.689 (0.668~ 0.739) | 0.644 (0.597~ 0.722) | 0.754 (0.720~ 0.787) | 0.586 (0.476~ 0.625) | 0.824 (0.794~ 0.852) | 0.744 (0.721~ 0.769) | 0.802 (0.767~ 0.85) | 0.750 (0.718~ 0.782) | 0.723 (0.697~ 0.845) |
| Sensitivity | 0.762 (0.647~ 0.800) | 0.741 (0.575~ 0.811) | 0.898 (0.845~ 0.936) | 0.557 (0.472~ 0.781) | 0.911 (0.850~ 0.942) | 0.874 (0.831~ 0.914) | 0.743 (0.592~ 0.809) | 0.801 (0.739~ 0.864) | 0.806 (0.582~ 0.854) |
| Specificity | 0.662 (0.636~ 0.767) | 0.606 (0.525~ 0.763) | 0.704 (0.663~ 0.754) | 0.607 (0.343~ 0.674) | 0.788 (0.750~ 0.843) | 0.716 (0.689~ 0.743) | 0.830 (0.769~ 0.960) | 0.736 (0.697~ 0.772) | 0.700 (0.669~, 0.919) |

Data enclosed in the parentheses represent the corresponding lower bound and the upper bound of 95% confidence interval. The intervals were obtained by bootstrapping strategy with 2000 random resamples.

Table S5

QUANTITATIVE RESULTS OF THE PROPOSED MODEL ON FFPE SLIDES AT TERTILE THRESHOLD.

| | BLCA | CESC | COAD | KIRP | LUAD | LUSC | SKCM | STAD | TNBC |
|---|---|---|---|---|---|---|---|---|---|
| AUC | 0.81 | 0.80 | 0.84 | 0.83 | 0.71 | 0.61 | 0.71 | 0.82 | 0.54 |
| Accuracy | 0.76 | 0.75 | 0.82 | 0.81 | 0.67 | 0.61 | 0.62 | 0.79 | 0.61 |
| Sensitivity | 0.78 | 0.77 | 0.67 | 0.67 | 0.72 | 0.54 | 0.82 | 0.70 | 0.54 |
| Specificity | 0.77 | 0.73 | 0.94 | 0.84 | 0.63 | 0.70 | 0.52 | 0.80 | 0.62 |

Table S6

Quantitative Results on FFPE Slides using The Model Trained with Fresh-frozen Slides.

| | BLCA | CESC | COAD | KIRP | LUAD | LUSC | SKCM | STAD | TNBC |
|---|---|---|---|---|---|---|---|---|---|
| AUC | 0.68 | 0.73 | 0.72 | 0.31 | 0.64 | 0.67 | 0.68 | 0.80 | 0.50 |
| Accuracy | 0.63 | 0.64 | 0.67 | 0.59 | 0.65 | 0.68 | 0.65 | 0.74 | 0.61 |
| Sensitivity | 0.79 | 0.89 | 0.80 | 0.31 | 0.70 | 0.63 | 0.73 | 0.76 | 0.38 |
| Specificity | 0.52 | 0.49 | 0.57 | 0.71 | 0.62 | 0.71 | 0.64 | 0.73 | 0.74 |

Table S7

HYPERPARAMETER SETTINGS OF COMPARISON METHODS

| Methods | Optimizer | Learning rate | Batch size | Patch size |
|---|---|---|---|---|
| Campella's | ADAM<br>$\beta_1$=0.9, $\beta_2$=0.999, $\epsilon$=1e-8 | 1e-4 | 512 | 256×256 |
| TransMIL | Rectified Adam<br>$\beta_1$=0.9, $\beta_2$=0.999, $\epsilon$=1e-8 | 2e-4 | 1 | 256×256 |
| CLAM | ADAM<br>$\beta_1$=0.9, $\beta_2$=0.999, $\epsilon$=1e-8 | 1e-4 | 1 | 256×256 |

Table S8

QUANTITATIVE RESULTS OF THE PROPOSED MODEL ON OTHER TUMORS AT THE THRESHOLD OF UPPER QUARTILE.

| | ACC | ESCA | HNSC | LIHC | MESO | OV | PRAD | READ | TCGT | THCA | UCEC |
|---|---|---|---|---|---|---|---|---|---|---|---|
| AUC | 0.819 | 0.496 | 0.719 | 0.558 | 0.707 | 0.644 | 0.629 | 0.828 | 0.763 | 0.838 | 0.641 |
| Accuracy | 0.707 | 0.560 | 0.630 | 0.637 | 0.677 | 0.584 | 0.501 | 0.680 | 0.811 | 0.779 | 0.526 |
| Sensitivity | 0.800 | 0.650 | 0.790 | 0.459 | 0.625 | 0.664 | 0.786 | 0.832 | 0.693 | 0.769 | 0.777 |
| Specificity | 0.685 | 0.514 | 0.584 | 0.692 | 0.697 | 0.544 | 0.417 | 0.651 | 0.850 | 0.783 | 0.439 |

Table S9

QUANTITATIVE RESULTS OF THE PROPOSED MODEL ON OTHER TUMORS AT THE THRESHOLD OF UPPER TERTILE.

| | ACC | ESCA | HNSC | LIHC | MESO | OV | PRAD | READ | TCGT | THCA | UCEC |
|---|---|---|---|---|---|---|---|---|---|---|---|
| AUC | 0.550 | 0.621 | 0.736 | 0.463 | 0.673 | 0.596 | 0.657 | 0.818 | 0.719 | 0.818 | 0.614 |
| Accuracy | 0.548 | 0.596 | 0.682 | 0.417 | 0.619 | 0.524 | 0.689 | 0.765 | 0.736 | 0.728 | 0.590 |
| Sensitivity | 0.733 | 0.787 | 0.830 | 0.934 | 0.738 | 0.881 | 0.471 | 0.750 | 0.632 | 0.831 | 0.662 |
| Specificity | 0.453 | 0.488 | 0.623 | 0.132 | 0.542 | 0.286 | 0.799 | 0.769 | 0.794 | 0.676 | 0.555 |

Table S10

QUANTITATIVE RESULTS OF THE PROPOSED MODEL ON OTHER TUMORS AT THE THRESHOLD OF MEDIAN.

| | ACC | ESCA | HNSC | LIHC | MESO | OV | PRAD | READ | TCGT | THCA | UCEC |
|---|---|---|---|---|---|---|---|---|---|---|---|
| AUC | 0.535 | 0.646 | 0.691 | 0.493 | 0.672 | 0.596 | 0.669 | 0.804 | 0.796 | 0.833 | 0.638 |
| Accuracy | 0.556 | 0.640 | 0.658 | 0.559 | 0.619 | 0.612 | 0.629 | 0.760 | 0.736 | 0.787 | 0.619 |
| Sensitivity | 0.541 | 0.374 | 0.545 | 0.583 | 0.736 | 0.696 | 0.816 | 0.808 | 0.563 | 0.813 | 0.739 |
| Specificity | 0.543 | 0.937 | 0.723 | 0.523 | 0.542 | 0.520 | 0.454 | 0.741 | 0.961 | 0.765 | 0.505 |

Table S11

PATIENT CHARACTERISTICS OF CPTAC COHORT

| Dataset | CPATAC-COAD (n=100) | CPTAC-BRCA (n=117) |
|---|---|---|
| Age (year) | | |
| Range | / | 31.3-91.3 |
| Average | / | 60.7 |
| Sex (n, %) | | |
| Female | 58 (58) | 106 (90.6) |
| Male | 42 (42) | / |
| Not reported | / | 11 (9.4) |
| Tumor Stage (n, %) | | |
| Stage I | 10 (10) | 4 (3.4) |
| Stage II | 41 (41) | 70 (59.8) |
| Stage III | 42 (42) | 32 (27.4) |
| Stage IV | 7 (7) | / |
| Not performed/reported | / | 11 (9.4) |

Table S12

Parameter Settings for Data Augmentation

| Methods | RandomVerticalFlip | RandomRotation | RandomResizedCrop | ColorJitter | Normalize |
|---|---|---|---|---|---|
| parameters | DEFAULT | Degree = (-90°, 90) | Size = (224, 224) | brightness=0.2, contrast=0.2, saturation=0.2, hue=0.2 | mean=(0.485, 0.456, 0.406) std=(0.229, 0.224, 0.225) |

Table S13

FITTED PARAMETERS OF CHI-SQUARE DISTRIBUTION

| Threshold | Degrees of freedom | Scale factor | Shifting |
|---|---|---|---|
| Upper quartile | 11.898 | 0.156 | -0.011 |
| Upper tertile | 6.794 | 0.234 | 0.178 |
| Median | 6.981 | 0.235 | 0.108 |
